# Sde Proteins Coordinate Ubiquitin Utilization and Phosphoribosylation to Establish and Maintain the *Legionella* Replication Vacuole

**DOI:** 10.1101/2023.09.07.553534

**Authors:** Kristin M. Kotewicz, Mengyun Zhang, Seongok Kim, Meghan S. Martin, Atish Roy Chowdhury, Albert Tai, Rebecca A. Scheck, Ralph R. Isberg

## Abstract

The *Legionella pneumophila* Sde family of translocated proteins promotes host tubular endoplasmic reticulum (ER) rearrangements that are tightly linked to phosphoribosyl-ubiquitin (pR-Ub) modification of Reticulon 4 (Rtn4). Sde proteins have two additional activities of unclear relevance to the infection process: K63 linkage-specific deubiquitination and phosphoribosyl modification of polyubiquitin (pR-Ub). We show here that the deubiquitination activity (DUB) stimulates ER rearrangements while pR-Ub protects the replication vacuole from cytosolic surveillance by autophagy. Loss of DUB activity is tightly linked to lowered pR-Ub modification of Rtn4, consistent with the DUB activity fueling the production of pR-Ub-Rtn4. In parallel, phosphoribosyl modification of polyUb, in a region of the protein known as the isoleucine patch, prevents binding by the autophagy adapter p62. An inability of Sde mutants to modify polyUb results in immediate p62 association, a critical precursor to autophagic attack. The ability of Sde WT to block p62 association decays quickly after bacterial infection, as predicted by the presence of previously characterized *L. pneumophila* effectors that inactivate Sde and remove polyUb. In sum, these results show that the accessory Sde activities act to stimulate ER rearrangements and protect from host innate immune sensing in a temporal fashion.

## Introduction

*Legionella pneumophila* is a facultative intracellular bacterium that is the causative agent of Legionnaire’s disease, a pneumonia of high lethality that primarily occurs in the immunocompromised and individuals with depressed lung function^1, 2^. Disease is usually associated with aspiration of droplets from contaminated water sources harboring amoebae that act as the environmental intracellular reservoir to support growth of the bacterium^3^. Once deposited within the lungs, productive infection involves growth of *L. pneumophila* within alveolar macrophages followed by attempted neutrophil clearing, which accumulates during productive pneumonic disease^4^. The ability of the bacterium to grow within macrophages is tightly linked to an environmental lifestyle that involves growth within amoebae, as establishing a niche within amoebal species provides the selective pressure for acquisition of genes that promote intracellular growth^5, 6^. Therefore, human disease promoted by the bacterium is a consequence of targeting host cell functions that are largely conserved from amoebae to man.

Intracellular growth of *L. pneumophila* requires construction of the *Legionella-* containing vacuole (LCV), which is derived from host membranes and is tightly associated with host endoplasmic reticulum and the early secretory apparatus^7, 8^. The LCV avoids interaction with the endocytic pathway including the highly antimicrobial lysosome^9^. Biogenesis of the bacterial compartment requires the function of the Icm/Dot type IV secretion system that acts as a syringe to deposit translocated bacterial effector proteins into the surrounding membrane as well as into the cytosol^10, 11^. Every *Legionella* species member encodes at least 300 of these effectors, which are remarkably poorly conserved, with only seven proteins found in all species^12, 13^. The biochemical activities of a large number of these proteins have been identified, many of which perform posttranslational modifications (PTMs) on host proteins, conferring activation, inhibition, altered regulation or acquisition of new functions on the target proteins^14^. The most notable consequences of these activities are to maintain the integrity of the LCV^15^, promote association with proteins involved in endoplasmic reticulum dynamics^16^, inactivate host components that drive association with the endocytic compartment ^17^, and inhibit protein synthesis^18^.

One set of Icm/Dot system translocated proteins that drives a unique morphological transformation of the LCV is the Sde family (SidE, SdeA, SdeB and SdeC)^19, 20, 21, 22^. The importance of this family is emphasized by the discovery that at least five other *L. pneumophila* translocated substrates regulate the dynamics of this family^23, 24, 25, 26, 27, 28^. As is true of many Icm/Dot substrates, each member is a composite of three enzymatic activities located on separate domains (Fig. 1A)^29, 30, 31^. A mono-ADPribosyltransferase (mART) domain drives NAD-dependent ADPribosylation of diverse ubiquitin species at Arginine 42 (ADPr-Ub)^20, 22^. In turn, a phosphodiesterase-like domain (PDE) can use ADPr-Ub to either promote phosphoribosyl linkage of Ub (pR-Ub) to substrates, or hydrolysis to generate free phosphoribosylated-Ub^21, 22^. The DUB domain is an accessory activity that preferentially deubiquitinates Lys63-linked polyubiquitin chains^32^. A number of potential mammalian substrates have been shown to be pR-Ub modified by Sde members on serine residues that are located in relatively unstructured regions of proteins^29, 30^. Work with model substrates indicates that tyrosine residues may also be targeted^33^. Although the number of Sde substrates is likely to be quite large given the structural preference for pR-Ub modification, modifications of endoplasmic reticulum protein Rtn4^22^ and Golgi-associated GRASP family members appear most closely tied to events involved in LCV biogenesis^27, 34^. In the absence of DUB activity, there is accumulation of polyUb about the replication vacuole, with unknown consequences on replication vacuole formation or intracellular growth^32^.

**Figure 1.**
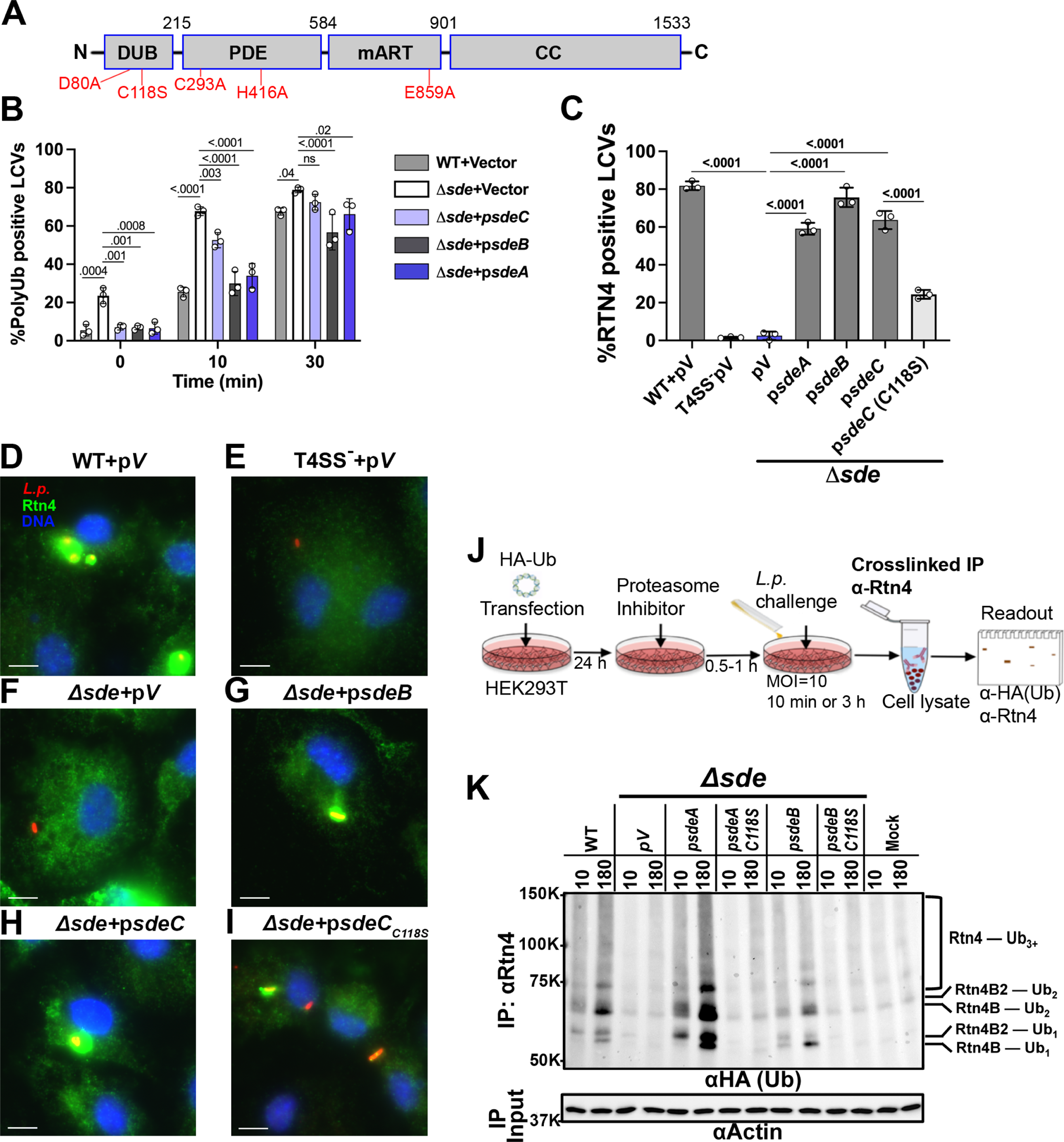
The Sde DUB domain is required for efficient phosphoribose-ubiquitination of Rtn4. **(A)** Domain structure of Sde family proteins. Catalytically inactive point mutations are shown in red. **(B)** Percent poly-Ub positive LCVs of *L. pneumophila* strains at noted timepoints post-infection. **(C)** Quantification of Rtn4-positive LCVs. BMDMs were challenged with indicated strains for the noted infection times, followed by fixation, permeabilization with 1% Triton X-100 and probing as described in Methods. 0 MPI was immediately after 5 min centrifugation of bacteria onto BMDM. Noted strains are wild type, Lp02; Δ*sde*, Δ*sidE*Δ*sdeC*Δ*sdeB-A*; Vector or pV, pJB908att-empty (Supplemental Table S1). For each experiment, n> 80 LCVs (B) or n>50 LCVs (C) per experiment were evaluated and data were determined from 3 biological replicates (mean ± SEM; two-way ANOVA with Tukey’s multiple comparison **(B, C)**; ns (non-significant); exact P values are displayed over data, and are found in Source Data File. For each replicate, n value is shown in Source Data File. **(D-I):** Representative micrographs of Rtn4 recruitment, scale bar = 5 µm. Scalebar determined for each image independently and pasted on each image. Macrophages were challenged with either WT containing pV or noted mutant strains for 1 hr, fixed, permeabilized and probed with anti-*L. pneumophila* (Alexa Fluor 594 secondary, red), anti-Rtn4 (Alexa Fluor 488 secondary, green), and Hoechst (nucleus, blue). **(J)** Cartoon of strategy to identify pR-Ub modified Rtn4. **(K)** Immunoblot image of anti-Rtn4 immunoprecipitates by immunoprobing with anti-HA. To right of immunoblots are Rtn4 isoforms and modification status. HEK293T cells were challenged with noted *L. pneumophila* strains for either 10 or 180 mins. Modified Rtn4 substrates were immunoprecipitated using crosslinked anti-Rtn4, fractionated on 7.5% SDS-PAGE, and probed with anti-HA. Lanes: WT, Lp02; vector, Δ*sde*; p*sdeA*, Δ*sde* expressing SdeA; p*sdeA C118S*, Δ*sde* expressing SdeA DUB mutant; p*sdeB*, Δ*sde* expressing SdeB; p*sdeB C118S*, Δ*sde* expressing SdeB DUB mutant; MOCK, uninfected. Masses are apparent molecular weights as noted in kDal. Source data are provided as a Source Data file.

Immediately after *L. pneumophila* is internalized by macrophages, pR-Ub modification of Rtn4 drives replication vacuole association with endoplasmic reticulum tubules. This modification results in highly detergent-resistant structures that appear to wall off the LCV from the host cell cytosol^22^, in a process that contributes to the protection of the LCV from attack by early endosomal components that degrade the vacuole^35^. Simultaneously, there is accumulation of polyubiquitinated targets in the vicinity of the LCV^36, 37, 38, 39, 40^, which appear to be targets of the Sde DUB domain^32, 41^. Phosphoribosyl modification of polyUb by Sde proteins stabilizes these structures, by blocking wholesale reversal of polyubiquitination by both bacterial and host DUBs^42^. Accumulation of polyUb potentially provides a target for the host cell autophagy pathway that could drive degradation of the LCV^43, 44, 45^. *L. pneumophila* appears to have several systems for blocking autophagic recognition of the LCV, including degradation of central components of the autophagy machinery by the Icm/Dot effector RavZ^46^, degradation of syntaxin 17 by effector Lpg1137^47^, reduction of sphingosine levels by LpSpl^48^ as well as the described phosphoribosylation^49^.

In this study, we report the role of the DUB and phosphoribosylation accessory activities of Sde family members. We show that the DUB domain surprisingly increases the efficiency of Rtn4 mobilization in the vicinity of the LCV, while ADPribosylation or phosphoribosylation of Ub interferes with autophagy by blocking binding of polyUb to an autophagy adaptor, primarily by phosphoribosylation of target Ub chains in the LCV vicinity. Similar results have been obtained in the simultaneously submitted manuscript by Wan, *et al.*^50^

## Results

The Sde DUB activity is required for efficient Rtn4 rearrangements in response to *L. pneumophila*.

To determine if there is a link between the Sde family K63 deubiquitinase (DUB) domain (Fig. 1A) and biogenesis of the *Legionella-*containing vacuole (LCV), the kinetics of polyubiquitin (polyUb) association and Rtn4 rearrangements in the vicinity of the LCVs were analyzed. A previous study had shown that the absence of the DUB domain resulted in a 2-fold increase in polyUb in the vicinity of the LCV^32^. Bone marrow-derived macrophages (BMDMs) were challenged with *L. pneumophila* derivatives lacking the complete complement of *sde* family members (*sidE*, *sdeABC*), which removes all proteins known to control Rtn4 dynamics^19^. After 5 min centrifugation onto BMDMs, followed by fixation and probing with a polyUb-specific antibody (FK1), there was no evidence of polyUb in the vicinity of the LCV of a WT strain containing empty vector (Fig. 1B; 0 minutes post infection (MPI)). In contrast, challenge with *L. pneumophila* missing all *sde* family members (Δ*sde*) resulted in approximately 24% of the LCVs displaying polyUb association. By 10 MPI, ∼25% of the vacuoles containing WT strain positively stained with polyUb, approximately 2-3X lower than the Δ*sde* strain at this timepoint, consistent with previous observations (Fig. 1B; ^32^). At 0 MPI, Δ*sde* mutants were efficiently complemented by all three members of the Sde family known to have biological activity (SdeA, SdeB and SdeC; Fig. 1B). Surprisingly, for all complemented strains, polyUb quickly rose to the levels seen in the absence of Sde family function (Fig. 1B; 30 MPI). The presence or absence of empty vector had no effect on the behavior of the WT strain (Supplemental Fig. S1). We conclude that inhibition of the polyUb association with the LCV occurs transiently. The fact that the inhibition is transient is consistent with the LCV-associated polyUb chains becoming resistant to the action of the DUB domain by 30 minutes post-infection (MPI).

The biological significance of having a K63 deubiquitinase activity is unclear, especially because the ART activity of Sde proteins directly blocks deubiquitination, arguing that the DUB may play some role other than reducing LCV polyubiquitination ^22, 42^. Alternatively, the DUB activity could modulate the dynamics of Rtn4 rearrangements associated with the LCV^22^. To test this model, BMDMs were challenged with *L. pneumophila* derivatives having defective DUB activity followed by probing for Rtn4 detergent-resistant structures in the LCV vicinity (Fig. 1C) ^22^. Surprisingly, the inactive DUB mutant (C118S) resulted in lowered Rtn4 colocalization with the LCV. In a strain expressing SdeC as the sole demonstrated driver of Rtn4 rearrangement, the C118S mutation reduced detergent-resistant Rtn4 association with the LCV by over 60% (Figs. 1C).

Consistent with data in Fig. 1C, Rtn4 association with the LCV was robust in strains expressing SdeB or SdeC as the sole plasmid-expressed isoforms (Figs. 1G,H) and indistinguishable from the WT strain (Fig. 1D). In contrast, loss of *sdeC-A* prevented Rtn4 association with the LCV, as did elimination of the T4SS (Figs. 1E, F). The presence of the C118S mutation resulted in two populations, with a mixture of detectable Rtn4-colocalization with LCVs or RTN4 staining below the level of detection (Fig. 1I). Similar results were seen with the DUB D80A catalytic mutant, whereas a control Cys mutant C293A showed high levels of colocalization (Supplemental Fig. S2). These results are consistent with the DUB domain playing a role in LCV biogenesis at the level of controlling Rtn4 dynamics.

To test if the Sde DUB activity plays a direct role in promoting Rtn4 rearrangements, we determined if the catalytic domain contributed to the rate of Sde-dependent phosphoribosyl-ubiquitin (pR-Ub) modification of Rtn4. In a purified system, pR-Ub modification of Rtn4 is rapid and robust with the presence of excess mono-Ub^22, 33^. In contrast, infection provides multiple potential sources of Ub, including K63-linked polyUb, which the DUB activity could target to provide substrate for pR-Ub modification. To test this hypothesis, HEK293T cells transfected with HA-Ub were challenged by *L. pneumophila* harboring either WT or DUB-defective (C118S) derivatives as the only active Sde family members. At 10 or 180 MPI, cells were extracted and subjected to immunoprecipitation with anti-Rtn4, followed by immunoprobing with anti-HA to reveal Rtn4 ubiquitination (Fig. 1J). Detectable mono-and di-modification of the Rtn4B and Rtn4B2 isoforms were observed as early as 10 MPI with continued increase at 180 MPI in response to the WT strain (Fig. 1J, WT). In contrast, challenge with Δ*sde* mutants showed HA-Ub modification that was indistinguishable from mock-infected controls (Fig. 1K; vector vs. Mock). Challenge with Δ*sde* mutants harboring single SdeA or SdeB isoforms each showed rescue of the HA-Ub modification defect, and their rescue was largely eliminated by the presence of the C118S catalytic mutation (Fig. 1K; compare p*sdeA* and *psdeB* to the respective C118S derivatives). Therefore, the absence of Sde DUB activity results in a strong reduction in both detergent-resistant Rtn4 and Ub modification of Rtn4, strongly arguing that Sde-mediated deubiquitination of K63-linked polyUb plays a role in LCV dynamics, possibly providing a pool of mono-Ub substrate for the ART domain.

### ADPribosylation of ubiquitin by Sde results in hyper-ubiquitination of the LCV

The presence of robust polyUb association with the LCV in the absence of the DUB domain indicates that pR-modification probably stabilizes polyUb (Fig. 1). This is inconsistent with the previous model that pR modification of Ub plays a role in blocking polyubiquitination during LCV formation^21^. Rather, it seems likely that pR-modification and ADP-ribosylation act to block both bacterial DUBs as well as a variety of host-encoded K63- and K48-specific DUBs (Fig. 2A)^42^. This was verified by incubating poly Ub_3-7_ (Ub chain lengths 3-7) with WT SdeC and active site mutants prior to incubation with either a low linkage-specificity DUB (USP2) or K63- and K48-specific DUBs of eukaryotic origin. Both WT enzyme (promoting pR modification) and the PDE(-) mutant (promoting ADPr modification) uniformly blocked DUB activity on either K48- or K63-linked Ub_3-7,_ similar to what was observed previously (Fig. 2A)^42^. As observed previously, the absence of the ART domain resulted in quantitative hydrolysis of the K63 polyUB chain by the SdeC DUB domain, leaving monomeric Ub that was poorly resolved after probing with a poly-UB specific antibody (Fig. 2A, ART^-^) ^22^. As predicted, this hydrolysis in the absence of ART activity was less efficient with the K48-linked substrate^22, 42^. The fact that ADPr modification blocked DUB activity allowed us to interrogate the consequences of Ub modification without confounding Rtn4 rearrangements promoted by the PDE domain.

**Figure 2.**
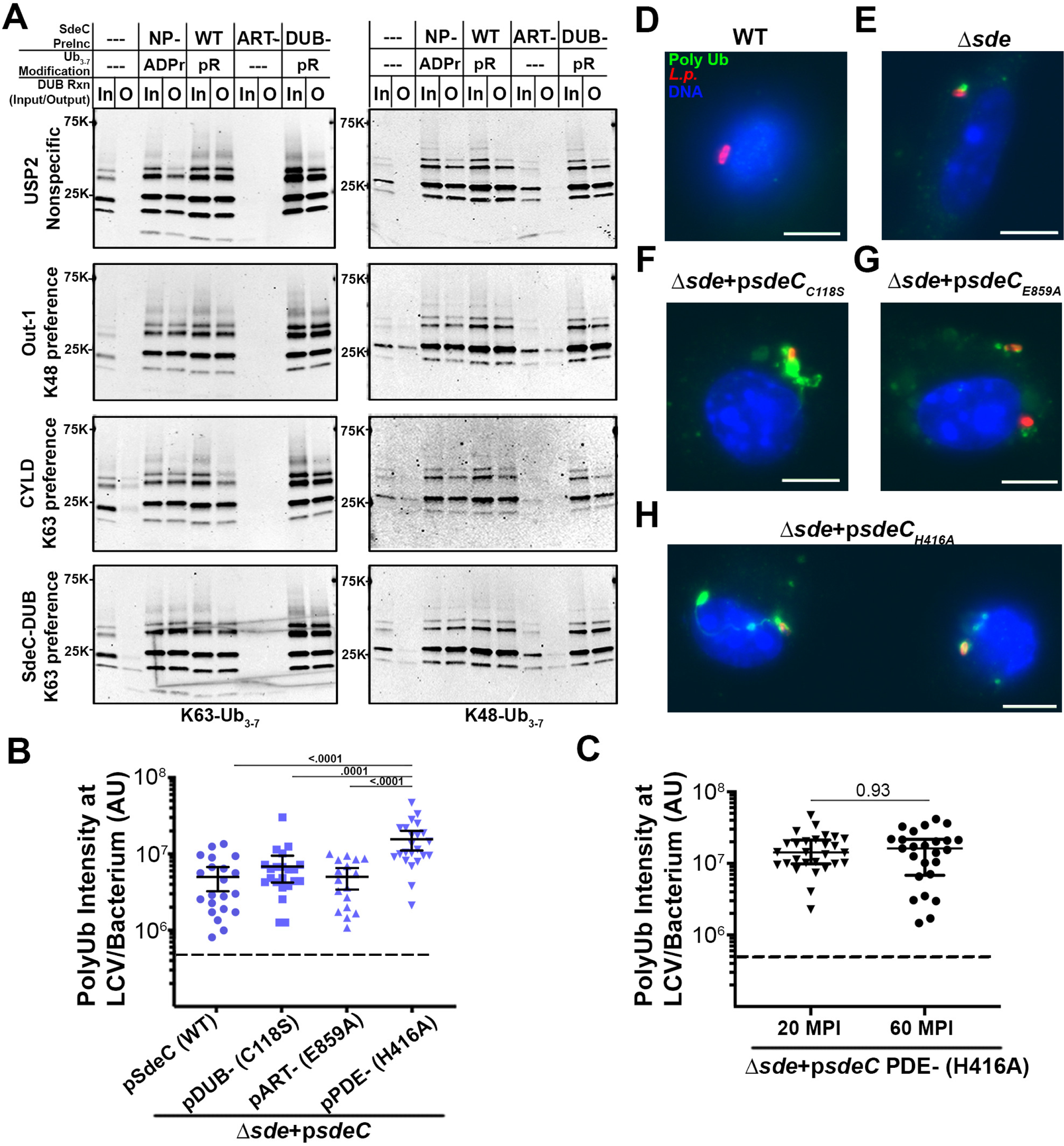
The ART domain blocks DUB activity and promotes hyper-polyubiquitination of the LCV. (**A**). SdeC WT or its catalytic mutant derivatives (1μg) were adsorbed to Ni-NTA resin for 1 hr at 4°C, then incubated with either K63 or K48-linked polyUb_3-7_ (2μg) for 2 hrs at 37°C. SdeC resin was removed and polyUb_3-7_ was then incubated with: human recombinant Polyhistidine-otubain 1, isoform 1 (100 nM); recombinant USP2 catalytic domain (50 nM); recombinant Polyhistidine-CYLD (50 nM); or recombinant SdeC DUB CD 1-192 (50 nM) at 37°C for 2 hrs. Cleavage was monitored by immunoblotting with anti-Ub (FK1). Shown are data from one of two independent replicates performed on separate days with similar results. Molecular weights are shown to left of each blot using protein standards of apparent molecular masses 25 kDal and 75 kDal. (**B**) Quantification of polyUb intensity associated with individual LCVs of the noted strains at 20 MPI. Data shown are means ± 95% confidence intervals, analyzed by one-way ANOVA with Dunnett’s multiple comparison. Data are pooled from 3 biological replicates. WT, n=21; C118S, n=23; E859, n= 17; H416, n=28. The dotted lines represent background level of polyUb intensity about LCVs**. (C)** PolyUb intensity of LCVs harboring Δ*sde* expressing SdeC PDE^-^ (H416A) mutant at 20 or 60 MPI. Data are shown in median ± 25%ile; Mann-Whitney 2-tailed test. The dotted lines represent background level of polyUb intensity about LCVs. Data are pooled from 3 biological replicates. 20 MPI, n=28; 60 MPI n=25. All P values are found in Source Data File. **(D-H)** Examples of polyUb-positive LCVs. Shown are images from one of three biological replicates with similar results. BMDMs were challenged by 5 min centrifugation at an MOI of 1, incubated with noted *L. pneumophila* strains, fixed, permeabilized with 1% Triton X100, and probed with anti-polyUb (Alexa Fluor 488 secondary, green), anti-*L. pneumophila* (Alexa Fluor 594 secondary, red), and Hoechst (blue). Strains used: WT, Lp02; *Δsde*, *ΔsidE ΔsdeC ΔsdeB-A; Δsde psdeC_WT_*, Δ*sde* expressing SdeC WT; *Δsde psdeC_C118S_*, Δ*sde* expressing SdeC DUB mutant; *Δsde psdeC_E859A_*, Δ*sde* expressing SdeC ART mutant; *Δsde psdeC_H416A_*, Δ*sde* expressing SdeC PDE mutant. Scale bar: 5μm. Source data are provided as a Source Data file.

To analyze *L. pneumophila* derivatives altered in Sde family function, BMDM monolayers were challenged for 20 min, and changes in polyUb in the LCV vicinity were analyzed by probing fixed samples with a polyUb-specific antibody (Fig. 2B). Neither *sdeC*_C118S_ (DUB^-^) nor *sdeC*_E859A_ (ART^-^) mutants showed significant increases in polyUb fluorescence intensity/vacuole if the analysis were restricted to LCVs having associated polyUb (Fig. 2B). In contrast, challenge with the *L. pneumophila* PDE^-^ mutant resulted in dense accumulation of LCV-associated polyubiquitination in spite of the fact that Rtn4 rearrangements were blocked (Fig. 2B). This dense association was maintained over time (Fig. 2C). Most notable was the presence of unusual polyUb-modified structures after challenge with the PDE^-^ mutant. While the Δ*sde* and ART^-^ mutants showed polyUb staining about individual LCVs (Figs. 2E,G), the PDE^-^ mutant showed polyUb structures that wrapped around the LCV and extended into serpentine structures across the interior of the BMDM (Fig. 2H). We occasionally observed dense structures in response to challenge with the DUB^-^ mutant as well (Fig. 2F), but the frequency was little changed from WT (Figs. 2B,C). We conclude that Sde-mediated modification of polyUb chains stabilizes LCV-associated Ub from the attack by both bacterial- and host derived DUBs. The observed deregulation of associated polyubiquitination in the absence of PDE activity indicates that direct modification of Ub is sufficient to stabilize polyUb. Furthermore, there is no evidence that pR- or ADPr-Ub modification plays a role in blocking polyubiquitination.

### Sde family members exclusively modify Arg42 residues throughout poly-Ub chains

The data showing that either pR- or ADPr-Ub modification stabilizes poly-Ub chains is consistent with Sde family members blocking recognition of poly-Ub by proteins that bind polyubiquitin chains. Previous work has shown that the Arg42 residue on mono-Ub is modified by either SdeA and SdeC^20, 21, 22^, indicating that phosphoribosylation of this residue is likely to be the critical event that blocks recognition by other proteins. However, no evidence has been provided that poly-Ub is quantitatively modified on the Arg42 residue, leaving the possibility that the blockade witnessed in our experiments may be due to alteration at other sites internal to poly-Ub. To demonstrate that internal Ub monomers within poly-Ub chains are quantitatively modified at Arg42, K63-linked Ub_4_ tetramers were incubated with SdeC-WT protein in the presence of NAD and analyzed by liquid chromatography-mass spectrometry (LC-MS; Fig. 3A; Methods)^22^. In the absence of SdeC, unmodified Ub_4_ protein was observed with a deconvoluted molecular mass of 34,206.75 AMU. Incubation of SdeC with Ub_4_ at a ratio of 1:750 (SdeC: Ub_4_) for 1 hr resulted in 96.5% of the Ub_4_ population increasing in mass by 848.21 AMU, corresponding to four modifications with mass=212.05 AMU, as predicted by four pR additions. These modified and unmodified Ub_4_ preparations were digested with trypsin, and subjected to LC-MS/MS to determine if the pR modifications were found exclusively on Arg42 (Fig. 3B), as they were for monomeric Ub^22^. An ion unique to the SdeC-treated preparation was identified at *m/z*=627.6392, *z*=3, as predicted for the 10-residue tryptic peptide containing the pR-modified Arg42 residue and as shown by extracted ion chromatogram (EIC). In contrast, no such peptide was present in the untreated samples (Fig. 3B). Furthermore, peptides containing the other Arg residues in Ub, Arg54 and Arg72, were indistinguishable in the SdeC-treated and untreated control, arguing against their modification in poly-Ub. A two amino acid tryptic fragment containing Arg74 was too small to be resolved in either sample.

**Figure 3.**
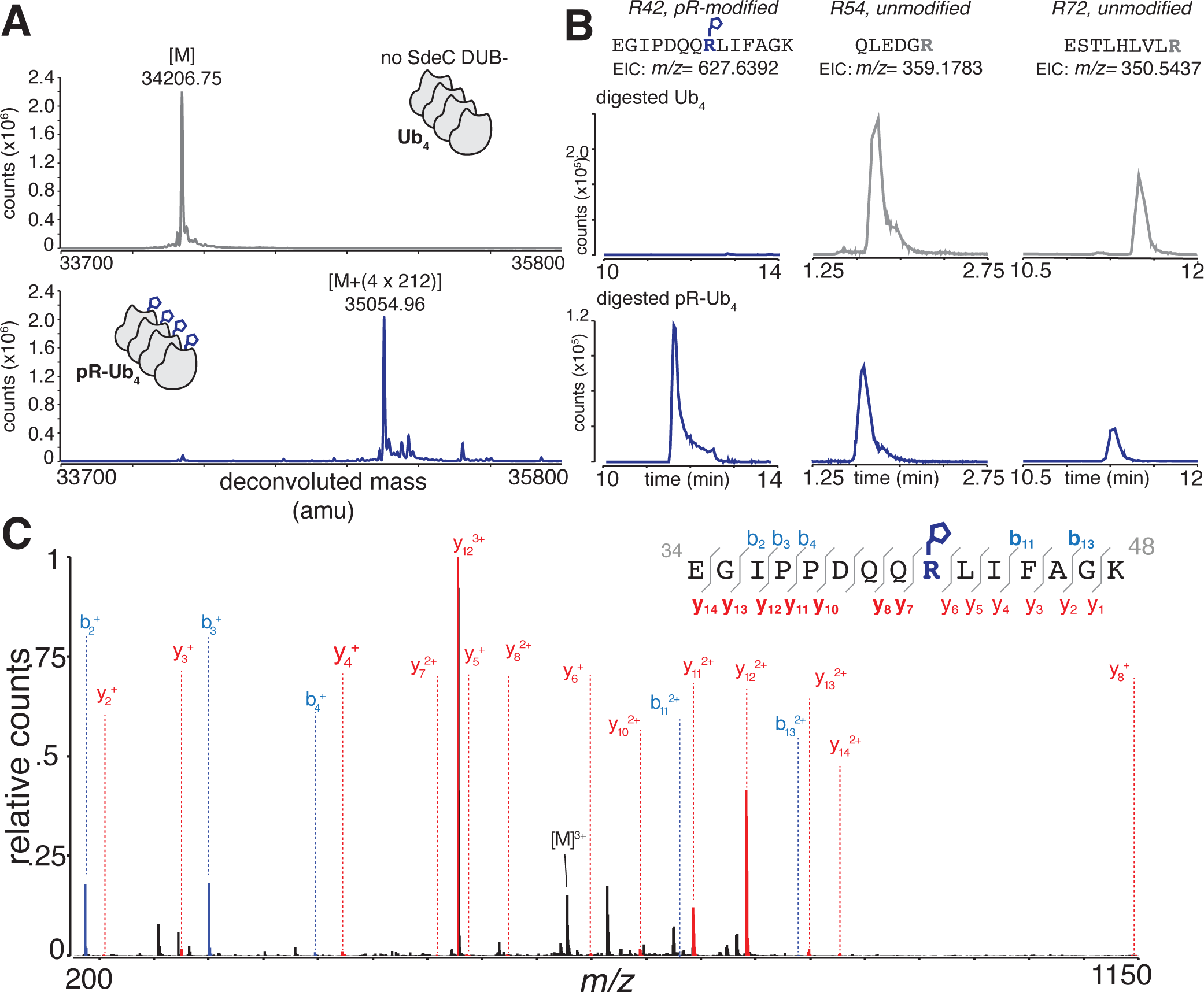
Phosphoribosyl modification of Ub_4_ occurs at Arg42. **(A).** SdeC(DUB^-^) incubation with Ub_4_ results in a mass increase of 848.21 AMU. Samples were subjected to LC-MS analysis and the deconvoluted masses of the peaks for each sample are displayed. Top: untreated Ub_4_. Bottom: Ub_4_ incubated with SdeC(DUB^-^) and NAD. **(B)**. Sde modification of Ub_4_ occurs exclusively on Arg42. Representative extracted ion chromatograms (EICs) are shown for the pR-modified R42 tryptic fragment, as well as tryptic fragments containing R54 and R72. The unmodified R42 fragment could not be found in SdeC-treated samples. (**C).** MS/MS analysis of the pR-modified tryptic fragment. Observed diagnostic ions that confirm specificity to R42 are shown in bold. Blue: b ions. Red: y ions. Source data are displayed. Underlying data are available n ProteomeXchange repository (www.proteomexchange.org/).

The 10-residue fragment from the Sde-treated sample was subjected to b- and y-ion analysis after LC-MS/MS to demonstrate that Arg42 was the modified residue (Fig. 3C). Modification of R42 by pR predicts that y ions beginning with y_7_ and successively larger peptide ions should have an increase in mass of 212.05 AMU, while y_6_ and shorter peptides should show no such increase. These ions were identified with high resolution, as were many other y ions leading up to, and beyond R42, yielding *m/z* ratios expected of pR modification at this site and not at other residues. Analysis of b ion data supported the y ion identifications. Additionally, no ions corresponding to the mass of unmodified R42 could be identified in SdeC-treated samples. Therefore, modification of poly-Ub occurs with the identical specificity of mono-Ub and similarly results in quantitative addition of pR to R42 using limiting amounts of enzyme relative to substrate.

### Sde-mediated modification of polyUb blocks autophagy adaptor SQSTM1/p62 binding via UBA domain

The Ub Arg42 residue is a key target for modulating ubiquitin dynamics. Autophagy adaptors such as p62 link ubiquitinated compartments to LC3-decorated phagophores via ubiquitin associated (UBA) domains that recognize a hydrophobic patch on the Ub surface^51, 52^. Integral to this patch are the residues I44, V70 and L71, directly proximal to R42, the target residue of ADPr modification by Sde (Fig. 4A, blue arrows). Structural analysis has shown that R42 can directly participate in the UBA interface (Fig. 4B) and is likely involved in p62 recognition based on NMR analysis^53, 54^. We therefore tested if modification of the Ub at Arg42 residue blocks recognition by the p62 autophagy adaptor.

**Figure 4.**
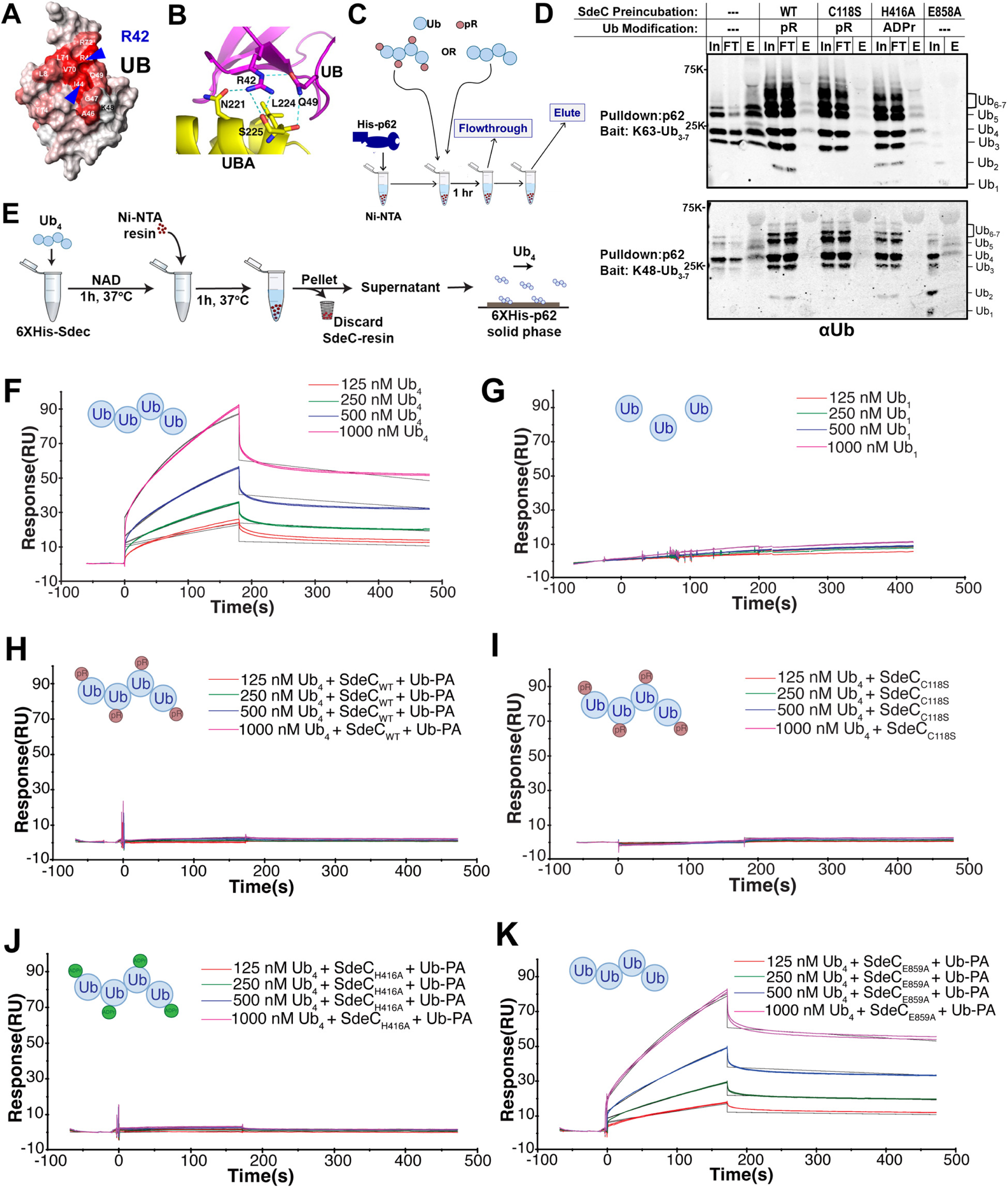
Sde-mediated modification of polyUb prevents its recognition by p62. **(A)** Space filling model of Ub. R42 residue (blue letter) and I44 residue are indicated with blue arrows (from reference^53, 59^).**(B)** Interface residues between UBA domain and Ub, showing possible involvement of Ub R42 in interacting with UBA domain (from reference^53, 59^). **(C, D)** Ub modification at R42 blocks p62 binding (Methods). Ni-NTA-p62 was incubated with Ub_3-7_ that had been treated with SdeC WT or its catalytic mutant derivatives and allowed to bind (In). Unbound Ub_3-7_ was removed (FT), the beads were washed, and then eluted with 250mM imidazole (E: elution). The p62-bound poly-Ub in the eluate was fractionated by SDS-PAGE followed by probing with anti-Ub. Relative loading: In, input (2.5%); FT, flow-through (2.5%); E, eluate (25%). Molecular weight markers are from protein standards of 25 kDal and 75 kDal apparent molecular weights. Shown are data from one of two independent experiments. (**E)** Surface plasmon resonance (SPR) quantitation of p62 binding to Ub derivatives. Ub_1_ or Ub_4_ was incubated with SdeC WT or its derivatives immobilized on Ni-NTA resin, the supernatant containing modified Ub derivatives was incubated with 6xHis-tagged p62 bound to a Ni-NTA chip, and binding affinity was measured by plasmon resonance. **(F, G)** Sensorgrams of Ub_4_ or Ub_1_ binding to p62. Binding of Ub_4_ to p62 yields a K_D_= 0.97 x 10^-7^ M. **(H-K)** Sensorgrams of Ub_4_, modified as noted in panels (red balls: phosphoribosyl addition; green balls: ADP-ribosyl addition). Reactions were formed in the presence of Ub-propargylamide (DUB inhibitor). All sensorgrams are source data. All other data are available as a Source Data file. Source data are provided as a Source Data file.

His-p62-Ni-NTA resin was challenged with K48- or K63-linked Ub_3-7_ mixed polymers that had been pretreated with SdeC WT or its catalytic mutant derivatives (Methods). The unbound fraction was collected (FT; flow through), and the p62-associated Ub_3-7_ (Eluate) was detected after co-elution with imidazole, followed by SDS-PAGE fractionation and immunoprobing with anti-Ub (Fig. 4C). In the absence of SdeC, both K48- and K63-linked Ub_3-7_ were efficiently co-eluted with p62 (Fig. 4D; no SdeC preincubation). In contrast, pretreatment with the SdeC WT, DUB^-^ (C118S) or PDE^-^ (H416A) derivatives all reduced K63- or K48-linked Ub_3-7_ binding to p62 (Fig. 4D; E lanes). This indicates that either pR- or ADPr-modification of polyUb blocks p62 recognition. In the case of the K48 derivative, the loss of the ART domain restored Ub_3-7_-p62 coelution, however it caused depolymerization of K48-linked Ub_3-7_ into monomeric Ub, making the effects of this mutant difficult to evaluate in this assay (Fig. 4D; E859A).

To determine the effects of Ub Arg42 modification in a more quantitative assay, binding affinities to Ub_4_ were determined using surface plasmon resonance (SPR) in the presence of a DUB inhibitor (Ub-propargylamide, Ub-PA) to prevent degradation of the Ub_4_ polymer (Supplemental Fig. S3). K63-linked Ub_4_ was modified by SdeC WT or its catalytic mutant derivatives in the presence of Ub-PA (Methods), the modified K63-linked Ub_4_ was introduced onto flow chips with immobilized p62, and binding was measured by SPR over time (Fig. 4E). In the absence of SdeC, Ub_4_ bound His-p62 with apparent K_D_ = 1.2 × 10^-7^ to 6.2 × 10^-8^ M, indicating that binding affinity was at least as efficient as previous SPR studies using diUb as a substrate (K_D_ = 9.3 × 10^-8^ M ^55^) (Fig. 4F). Binding of p62 to mono-Ub was very poor, consistent with DUB activity interfering with recognition of p62, with binding constants that could not be reliably determined by SPR (Fig. 4G, Fig. S3). Ub_4_ pretreated with SdeC, either in the presence of Ub-PA or having the C118S DUB mutation, resulted in blockade of p62 binding, consistent with phosphoribosylation blocking recognition of Ub by UBA domains (Figs. 4H,I). ADPr modification of Ub at Arg42 was sufficient to block binding, as the SdeC_H416A_ PDE^-^ mutant blocked binding as efficiently as WT (Fig. 4J). In all cases in which modification blocked p62 recognition, the binding constants could not be reliably determined by SPR. Elimination of the ART activity restored Ub_4_ binding to p62, with a similar binding affinity to untreated, consistent with PA blocking the DUB activity (Fig. 4K; K_D_ = 1.6 × 10^-7^ M). These results strongly argue that modification of the Ub at Arg42 residue by SdeC disrupts recognition by p62.

### Sde-mediated modification of polyUb chains prevents association of p62 with the LCV

To determine if Sde proteins enzymatically camouflage the replication vacuole from recognition by p62, BMDMs were challenged with *L. pneumophila sde* variants and probed for p62 localization in the vicinity of the LCV. A variety of p62 morphological variants were observed, including LCV-associated puncta and LCVs enveloped by p62 (Supplemental Fig. S4). After bacterial contact (5 min centrifugation) and immediate fixation, LCVs harboring *L. pneumophila* WT showed no envelopment of p62 (Fig. 5A). In contrast, a large fraction of the vacuoles after challenge with the *L. pneumophila* Δ*sde* strain were enveloped by p62 (Fig. 5B). As time progressed, a variety of morphologies became apparent, such as the alignment of puncta proximal to the LCV. As puncta associated with the LCV were difficult to distinguish from those in the general vicinity, we limited quantification to identifying vacuoles with enveloping p62 structures and determining total p62 intensity associated with individual vacuoles.

**Figure 5.**
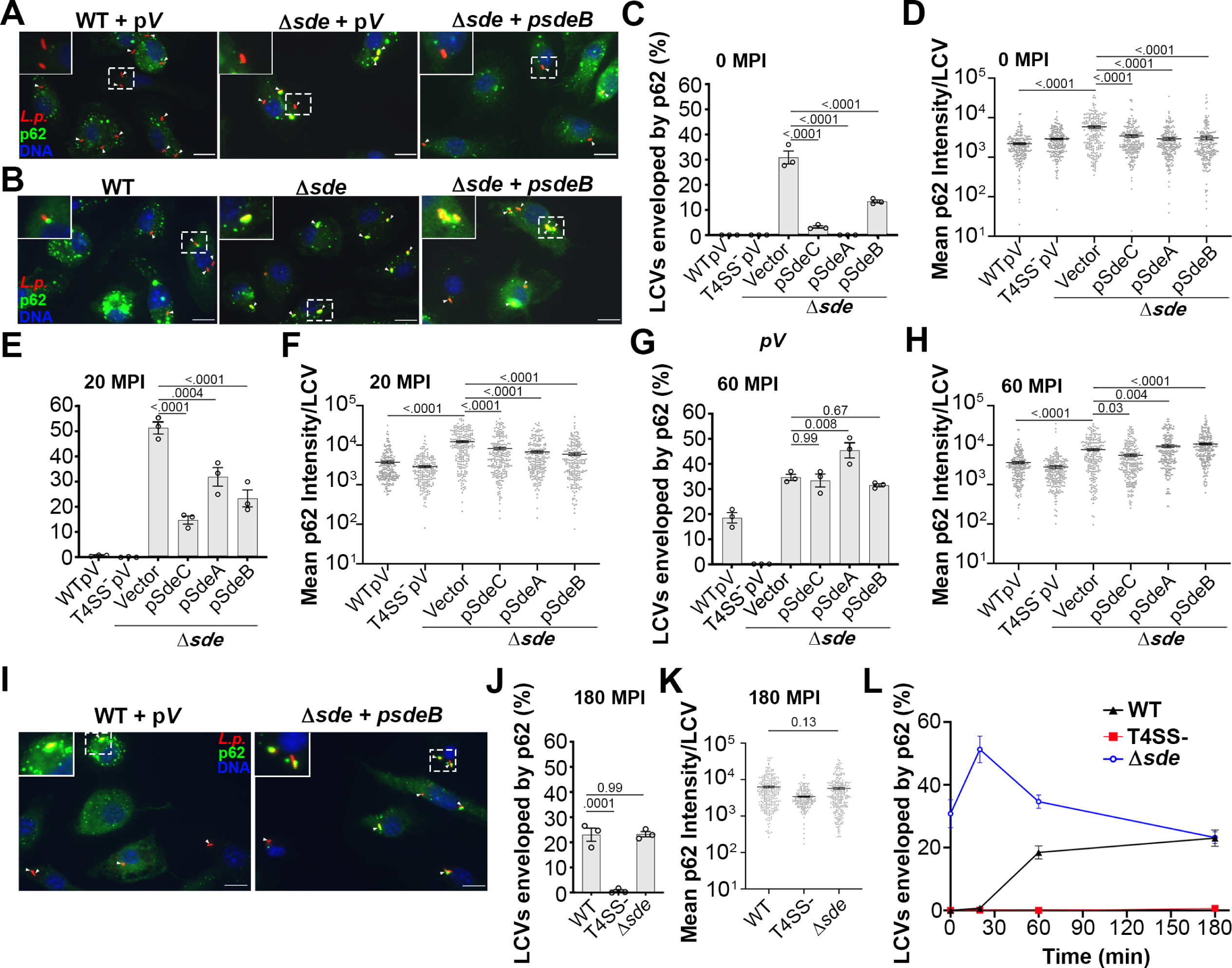
Transient blockade of p62 recruitment to the LCV in the presence of Sde family members. **(A-B)** Examples of punctate (A) or enveloped (B) p62 morphology localized around LCVs after challenge of BMDMs with noted *L. pneumophila* strains. The scale bar represents 10 µm. **(C, E, G, J)** The Percentage of circumferential p62 associated with LCVs at 0 (immediate bacterial contact; C) or 20 (E) or 60 (G) or 180 (J) MPI. 100 LCVs per replicate were counted in n=3 biological replicates (mean ± SEM; one-way ANOVA with Dunnett’s multiple comparison; ns (non-significant). P values are displayed over data. **(D, F, H, K)** Quantification of mean p62 intensity/pixel around LCVs at 0 (D) or 20 (F) or 60 (H) or 180 (K) MPI of vacuoles analyzed in (C,E,G,J). Image analysis performed on same coverslips at corresponding timepoints. Number of cells analyzed (n) for each strain at each timepoint is provided in Source Data File. At least 70 vacuoles per replicate were quantified and data were pooled from 3 biological replicates (mean ± SD; one-way ANOVA with Dunnett’s multiple comparison (D, F, H) and Student’s two tailed t-test (K); ns (non-significant). Exact P values are displayed over data. (**I**) A representative micrograph of p62 association with LCVs at 180 MPI, scale bar 10 µm. (**L**) Kinetics of blockade against p62 recruitment. All strains described in Supplemental Table S1 (pV: pJB908att-empty). Data acquired from panels C, E, G and J, and shown as mean ± SEM. All P values are found in Source Data File. Source data are provided as a Source Data file.

Immediately after bacterial contact with BMDMs, both the WT and the T4SS^-^ strains were devoid of p62 staining (Fig. 5C). In contrast, approximately 30% of the *L. pneumophila* Δ*sde* bacteria were found to be enveloped by p62 (Fig. 5C). Colocalization was largely lost when either *sdeA, sdeB, or sdeC* expressed on plasmids was introduced into the Δ*sde* strain. Quantification of p62 signal associated with the LCV showed that there were two populations of bacteria after challenge with the Δ*sde* strain, with a population showing p62 recruitment levels that were similar to WT and a second population with 8-10X higher fluorescence intensity (Fig. 5D). In contrast, the WT and Δ*sde* strains harboring Sde isoforms *in trans* showed indistinguishable low levels of p62 recruitment (Fig. 5D).

As the infection proceeded, colocalization of p62 with the Δ*sde* strain continued to increase (Fig. 5E, >50% at 20 MPI) with the population of LCVs strongly skewing toward hyperaccumulation of p62 (Fig. 5F). Concurrently, a second phenomenon became apparent in strains harboring plasmids that produced excess Sde isoforms. There was clear complementation of the Δ*sde* defect (Fig. 5E), but accumulation of p62 in the vicinity of the LCV at 20 MPI was observed that distinguished these constructs from the WT strain (Fig. 5E). Accumulation could also be observed when the fluorescence intensity of individual LCVs was determined, with a population emerging with increased p62 density (Fig. 5F). This phenomenon appeared to be a premature version of what occurs with the WT strain, as 20% of the LCVs harboring the WT showed accumulation of p62 by 60 MPI, with evidence of an emerging population showing increased p62 density in the LCV vicinity (Figs. 5G, H). There was robust accumulation of p62 about individual LCVs harboring WT strain at this timepoint that appeared indistinguishable from individual LCVs harboring the Δ*sde* strain (Fig. 5I). Furthermore, at 180MPI, accumulation of p62 around the LCV in the Δ*sde* strain was indistinguishable from p62 accumulation around WT LCVs (Figs. 5J, K). When the percentage of LCVs associated with p62 was plotted as a function of time post-infection (Fig. 5L), there was early accumulation of p62 that was blocked by the Sde family (0-20 MPI). This was followed by decay of p62 accumulation after infection of the Δ*sde* strain, while the WT showed increasing p62 between 20 and 60 MPI, resulting in a convergence of the phenotypes (180 MPI). The loss of Sde protection of the LCV is consistent with reversal of its activity by known metaeffectors SidJ and DupA/B ^24, 25, 26, 56^.

To determine the role of the individual Sde activities in preventing p62 accumulation, point mutations in each of the three catalytic domains were analyzed at 0 MPI after macrophage challenge, when the Sde family exerts its most striking effects. Mutations in *sdeB* were introduced into the Δ*sde* strain, as this isoform showed the most effective complementation (Fig. 5C), and these mutants were then used to challenge macrophages. Point mutations in the catalytic sites of the DUB or PDE domains, as well as a double mutant missing the catalytic activity in both of these domains, still retained the ability to block p62 colocalization after association (Figs. 6A-C). In contrast, a mutation in the ART domain caused a large increase in p62 recruitment to LCVs, indicating that ADP ribosylation of Ub Arg42 by the ART domain blocks p62 recognition of the LCVs (Fig. 6A). The presence or absence of empty vector in the WT strain had little effect on this result (Supplemental Fig. S5). It should be noted that there appeared to be some residual interference of p62 in the single ART mutant, but this was eliminated by simultaneously introducing a DUB mutation, indicating that the DUB domain may play a minor role in lowering p62 recognition of the LCV (Fig. 6A). As seen with the Δ*sde* strain, the ART mutants resulted in a clear subpopulation showing approximately 10-fold increase in p62 accumulations (Figs. 6B,C). These results support the biochemical data that either ADPribosylation or phosphoribosylation of Ub at Arg42 is sufficient to block p62 recognition. It also argues against models that propose pR-linked Ub modification of a peripheral factor plays a role^57^, as ADPribosylation of Ub was sufficient to interfere with p62 recruitment.

**Figure 6.**
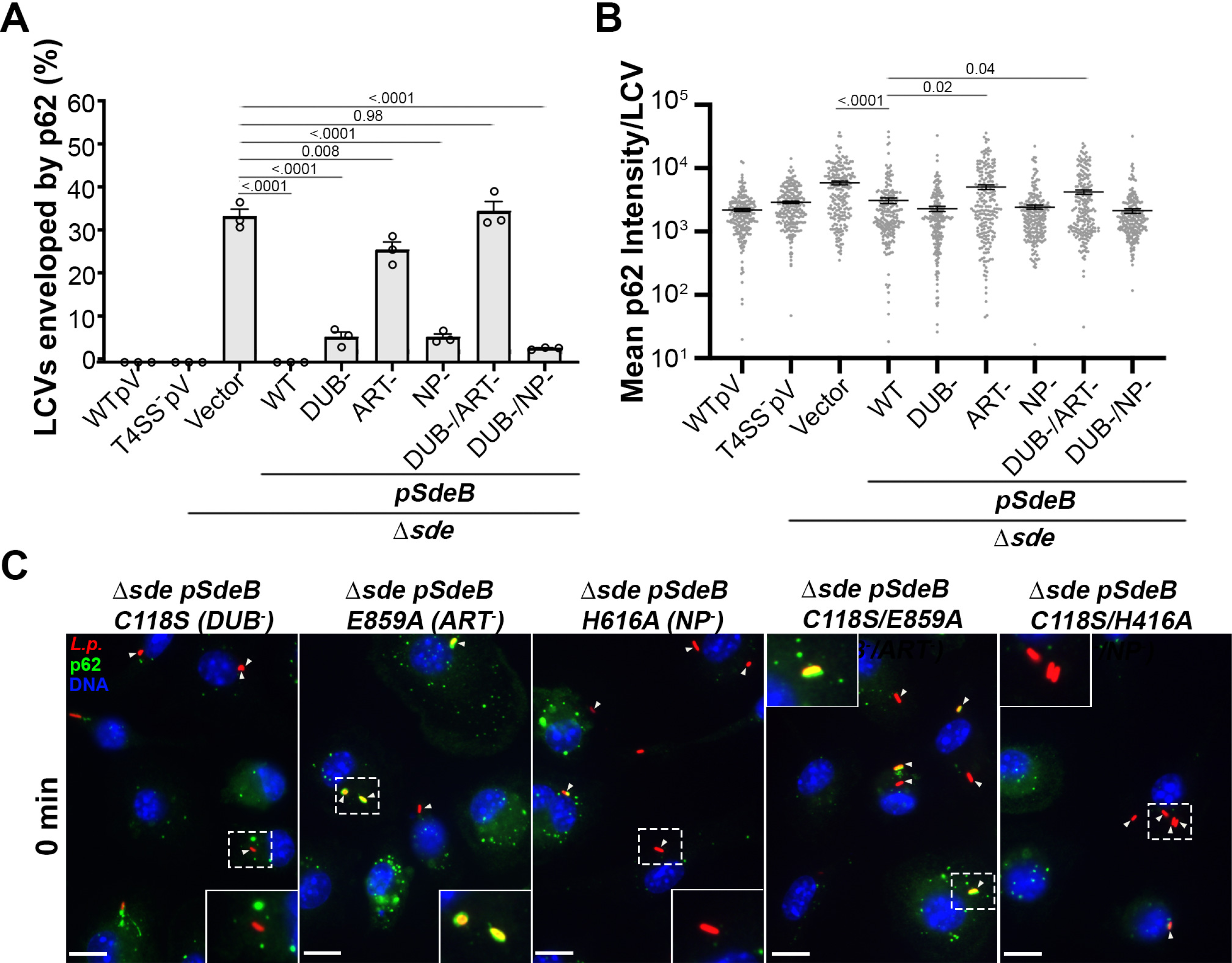
ART activity is sufficient to block p62 recruitment to the replication vacuole. **(A)** Quantification of individual LCVs enveloped by p62 at 0 MPI. BMDMs were challenged with the noted *Legionella* strains. 100 LCVs per replicate were counted in n=3 biological replicates (mean ± SEM; one-way ANOVA with Dunnett’s multiple comparisons; ns (non-significant). Exact P values are displayed over data. **(B)** Quantitation of mean p62 intensity/pixel associated with individual LCVs. Images of individual vacuoles were grabbed and pixel intensities of p62 staining about regions of interest were determined (Methods). At least 198 LCVs were counted from the replicates performed in panel A. Data are shown as means ± SD (one-way ANOVA with Dunnett’s multiple comparison; ns (non-significant). Exact P values are displayed over data. All P values are found in Source Data File. **(C)** Examples of recruited p62 to LCVs harboring Δ*sde* expressing SdeC catalytic mutant derivatives. All strains described in Supplemental Table S1 (pV: pJB908att-empty). Scale bar: 10 μm. Source data are provided as a Source Data file.

## Discussion

The *L. pneumophila* Sde family drives phosphoribosyl-linked ubiquitination (pR-Ub) of targets, most spectacularly leading to largescale rearrangements of the tubular ER protein reticulon 4 (Rtn4)^22^. In addition, there are two accessory activities of Sde proteins that have unclear biological roles. The Sde deubiquitinase (DUB) domain preferentially deubiquitinates Lys63-linked polyubiquitin chains (Fig. 1)^32^, while the other is the hydrolytic product of ADPribosylated ubiquitin (ADPr-Ub), resulting in phosphoribosylated ubiquitin (pR-Ub)^22, 58^. The phosphoribosylation occurs at the Ub Arg42 residue, strategically placed in a hydrophobic patch that provides a recognition surface for ubiquitin binding proteins that drive autophagy (Fig. 3C; Fig. 4A,B)^53, 59^. The presence of Sde family members is linked to a lowered association of Ub-recognizing autophagy adaptors to the LCVs^49^, consistent with phosphoribosylation promoting maintenance of the LCV in the face of cytosolic antimicrobial attacks.

In this manuscript, we showed that inhibition of polyUb formation about the replication vacuole was transiently controlled by the Sde DUB domain and limited to the earliest times after infection (Fig. 1). This early degradation of polyUb by the DUB domain fueled phosphoribosyl-linked ubiquitination because pR-Ub modification of Rtn4 slowed greatly in the absence of DUB activity and was accompanied by a reduction in Rtn4 accumulation about the replication vacuole (Fig. 1D-M). Therefore, it appears that there is K63-polyubiquitination of the LCV about the nascent replication vacuoles^39, 60, 61, 62, 63, 64^ that provides the substrate for the Sde DUB activity, generating monoubiquitin fuel for the Sde mART domain (Fig. 7B,C). This is then directly transferred by the PDE domain to generate pR-Ub modified Rtn4, allowing a direct pathway from vacuolar polyUb to endoplasmic reticulum (ER) rearrangements (Fig. 7D).

**Figure 7.**
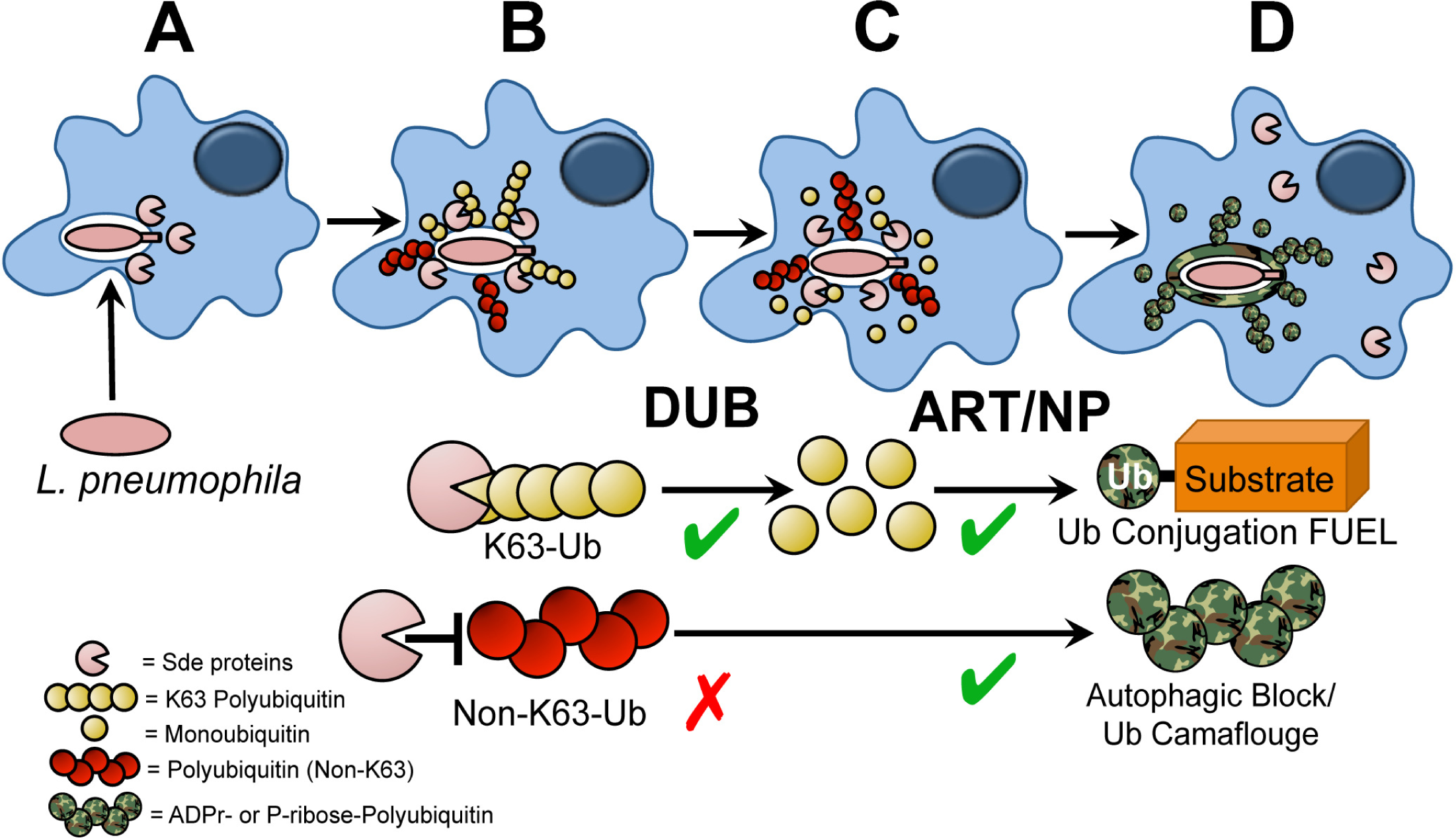
Sde accessory activities drive Rtn4 rearrangements and block autophagic recognition. The DUB activity provides fuel for pR-Ub modification of Rtn4, driving rapid ER-associated rearrangements. **(A)** Bacteria are internalized, Sde family proteins are translocated. **(B)** Polyubiquitination occurs about the replication vacuole. **(C)** Digestion of polyUb provides Ub fuel for ART/PDE domain to catalyze pR-Ub conjugation to substrates such as Rtn4. **(D)** Rtn4-pR-Ub modified protein accumulates about the replication vacuole polyUb is modified by phosphoribosylation. The pR-modification of the polyUb at R42 residue stabilizes polyubiquitination about LCVs, simultaneously preventing degradation by host cell-derived DUBs and blocking recognition by the autophagic machinery.

The association of polyUb with the replication vacuole should be a suicide signal for *L. pneumophila*, as ubiquitination marks intracellular compartments for degradation by the autophagy pathway^45, 65^. Autophagy results from binding of adaptor molecules (such as p62) to Ub-modfied compartments which, in turn, bridge to autophagophores decorated by LC3/Atg8^66^. It is well established that after *L. pneumophila* forms the LCV, autophagy is blocked via the action of the Icm/Dot substrate RavZ, which irreversibly hydrolyzes and deconjugates LC3/Atg8 from the nascent autophagophore, blocking binding to autophagy adaptors^46^. In the absence of RavZ, the LCV is still resistant to autophagy, indicating that the vacuole is masked in a fashion that prevents recognition by the LC3-decorated autophagophore^46, 49^. We show here that modifying the Arg42 residue of polyUb with either mono-ADPribose or phosphoribose physically blocks binding to the autophagy adaptor p62 (Fig. 4), and similar results have been obtained by Wan and coworkers^50^. Surface plasmon resonance experiments showed no detectable p62 binding in presence of high concentrations of polyUb modified by either WT (pR modification) or the H416A derivative (ADPr modification; Fig. 4F-K). Consistent with the data using purified proteins, both modifications blocked p62 association with the LCV (Fig. 6) and stabilized polyUb about the replication vacuole (Fig. 2). In fact, the ADPr modification resulted in hyperaccumulation of polyUb about the replication vacuole (Fig. 2B,C), consistent with enhanced resistance to both DUBs and autophagy adaptor recognition.

That all the proposed functions of phosphoribosylation can be fulfilled without hydrolysis of the ADPr modification questions the purpose of the pR modification. It seems likely that allowing vacuole-associated polyUb to accumulate ADPr modifications would result in covalent linkage of Sde targets directly to the replication vacuole, even if this is a low likelihood event ^22, 27, 34^. In addition, the ART and PDE domains work independently and consecutively, allowing ADPr modification to persist on polyubiquitin chains, raising the possibility that unwanted proteins would be immobilized about the replication vacuole. This problem is particularly exacerbated by the fact that Sde-promoted pR-Ub linkage preferentially occurs on unstructured regions of proteins, which could drive promiscuous linkage of poorly folded proteins about the vacuole. Therefore, hydrolysis of ADPr-Ub at Arg42 acts as a timing device, requiring target to be immediately accessible to Sde and ADPr-Ub prior to decay to pR-Ub.

It has been previously reported that pR-Ub modification of USP14, a host cell deubiquitinase, is a central strategy for preventing p62 association with the *Legionella* replication vacuole^57^. The authors proposed that modification of USP14 disrupted its direct interaction with p62, preventing association of the LCV with the autophagy adaptor^57^. The data we presented here argue against USP14 modification being a major mechanism for autophagy avoidance, and that modification of LCV-associated Ub is likely the primary strategy used by Sde proteins to protect against autophagy. First, pR-modification of Ub blocked p62 association (Fig. 4), and, given the expected stoichiometry of Ub relative to USP14, it is likely that this direct blockade is the primary contributor to autophagy avoidance. Secondly, ADPr-modification of poly-Ub was sufficient to avoid autophagy (Figs. 2B, 5A), arguing that pr-Ub modifications of any peripheral targets such as USP14 are unlikely to contribute to autophagy avoidance. Finally, we showed that there was a minor, but detectable, contribution of the DUB domain to autophagy avoidance (Fig. 5A), consistent with host DUBs making similar enzymatic contributions, rather than playing a role in direct binding to the p62 target.

Our data on the transient nature of p62 blockade illustrates an important biological principal underlying Sde family function (Fig 4L). There is little, if any, association of p62 with the *L. pneumophila* WT replication vacuole during the first 20 minutes of macrophage infection. In contrast, challenge with the Δ*sde* strain results in p62 association with the LCV at early timepoints (Figs. 4E,L). This huge difference largely disappears by 1 hour post infection, as there appears to be increased accumulation of p62 about the WT LCV, with coordinate decreased p62 accumulation about the Δ*sde* LCV (Fig. 4L). By 3 hr post infection, the Δ*sde* mutant and WT look indistinguishable, with each showing about 80% of the LCVs devoid of p62 localization. Therefore, function of Sde decays rapidly during the first hour after infection, with a new activity appearing in both strain backgrounds that interferes with p62 association.

The decay in protection from p62 association is predicted by the function of proteins that modulate Sde family activity. Icm/Dot translocated proteins SidJ and SdjA glutamylate Sde proteins, blocking their ability to ADPribosylate ubiquitin^23, 24, 25, 26^. As expression of the *sde* genes appears to be downregulated after contact with mammalian cells^19, 67^, this modification effectively removes the pool of enzyme capable of ADPribosylating Ub. The fact that over time there is some increase in the level of p62 association with the WT LCV indicates that there is accumulation of polyubiquitin in the LCV vicinity during the first hour after infection that cannot be modified by the inactivated Sde enzymes. In this regard, it should be noted that pR-Ub modification of target proteins is also reversed by the linkage-specific deubiquitinases DupA and DupB, consistent with a general decay in Sde function as well as loss of memory of its activity during the first hour after infection^27, 28^. Therefore, Sde functions as critical cog in the morphogenesis of the LCV during the earliest stages of replication vacuole establishment. Efficient intracellular growth requires decay in Sde function, as SidJ mutants that result in extended Sde family activity show slowed intracellular growth^68^.

In summation, the Sde family is involved in critical early steps that are associated with homeostatic regulation of replication vacuole morphogenesis. Accessory activities that modulate polyubiquitin dynamics about the replication vacuole are tightly linked to phosphoribosyl-linked ubiquitination of substrate (Fig. 7). We argue that the DUB domain provides fuel for pR-Ub modification, while pR modification of polyubiquitination associated with the LCV prevents attack by an important arm of host cell cytosolic defense. Finally, the transient nature of Sde function during replication vacuole formation is an intrinsic feature of this system, as we witness decay in the ability to block the autophagic response. The transient nature of providing fuel for Sde function is also supported by this work, as phosphoribosyl modification of polyUb is predicted to block DUB function (Fig. 2) and reduce the amount of fuel to support Sde-mediated ubiquitination of substrates. The biological importance of limiting Sde function to the earliest stages of replication compartment biogenesis requires further investigation.

## Methods

### Ethics statement

All research complies with biosafety and animal welfare regulations as overseen by the Tufts University Institutional Biosafety Committee and Institutional Animal Care and Usage Committee.

### Bacterial culture and media

*L. pneumophila* derivatives used in these studies were streptomycin-resistant restriction-defective thymidine auxotrophs derived from a clinical isolate of *Legionella pneumophila*, strain Philadelphia-1 (Lp01)^11, 69, 70^. *L. pneumophila* strains were propagated on charcoal–N-(2- acetamido)-2-aminoethanesulfonic acid (ACES)–yeast extract plates, with or without thymidine as needed (CYE/T) and in ACES-yeast extract broth (AYE/T)^7, 11, 71^. Plasmids were introduced as described^22^.

For the infection of mammalian cells, *L. pneumophila* strains were grown overnight to post-exponential phase (OD_600_ 3.5-4.5) to a predominantly motile state^72^. *Legionella pneumophila* derivatives are described in Supplemental Table S1, *Escherichia coli* derivatives are described in Supplemental Table S2 and oligonucleotides used in this study are listed in Supplemental Table S3.

### Eukaryotic cell culture

Mice were housed at the Tufts University School of Medicine IACUC facility, with housing and all manipulations detailed in approved protocol B2021-119. Bone marrow-derived macrophages (BMDMs) were isolated from femurs and tibias of female 6-10week old A/J mice^7^ and frozen in fetal bovine serum (FBS) with 10% dimethyl sulfoxide (DMSO). Mouse strain verifications provided by Jackson Laboratories (Bar Harbor, ME). Approximately 3 mice were sufficient for all experiments performed. Cells were plated the day before infection in RPMI 1640 (Gibco) supplemented with 10% FBS and 2 mM glutamine at 37°C with 5% CO_2_. HEK293T cells, obtained from ATCC (not independently authenticated after being obtained), were cultivated in Dulbecco modified Eagle medium (Gibco) supplemented with 10% heat-inactivated (FBS, Invitrogen) (vol/vol), 100U/ml of penicillin-streptomycin (Gibco) at 37°C with 5% CO_2_.

### Polyubiquitin colocalization with LCV

BMDMs were plated on glass coverslips at 1-2 x 10^5^/well in 24-well plates. Cells were challenged at a multiplicity of infection (MOI) = 1, centrifuged at 200 x g for 5 min and incubated for the indicated time points. The infection mixtures were then fixed in PBS containing 4% paraformaldehyde, permeabilized in PBS containing 1% (vol/vol) Triton X-100 for 30 min, probed with mouse anti-polyUb (Enzo Life Biosciences, Cat# BML-PW8805-0500, 1:250) and rabbit anti-*L. pneumophila* followed by secondary probing with goat anti-rabbit Alexa Fluor 594 (Invitrogen, Cat# 111-585-144, 1:500) and donkey anti-mouse Alexa Fluor 488 (Jackson ImmunoResearch, Cat# A-11012, 1:500). Hoescht 33342 (Life Technologies, Cat#62249 1:10,000) was used to label nuclei.

### Immunofluorescence of Rtn4-LCV colocalization

BMDMs were plated on glass coverslips at 1-2 x 10^5^/well in 24-well plates. Cells were infected at a multiplicity of infection (MOI) = 1, centrifuged at 200 x g for 5 min and incubated for 1 hr. The infection mixtures were then fixed in PBS containing 4% paraformaldehyde, permeabilized in PBS containing 1% (vol/vol) Triton X-100 for 30 min, probed rabbit anti-RTN4 (Lifespan Biosciences, Cat# LS-B6516, 1:500) and rat anti-*L. pneumophila*^73^ followed by secondary probing with goat anti-rabbit Alexa Fluor 488 (Jackson ImmunoResearch, Cat# 111- 545-003, 1:500) and donkey anti-rat Alexa Fluor 594 (Jackson ImmunoResearch, Cat# 712-585-153, 1:500). Hoescht 33342 (Life Technologies, Cat#62249 1:10,000) was used to label nuclei.

### Rtn4 Immunoprecipitations

For transfections preceding Rtn4 Immunoprecipitations (IP), HEK293T cells were plated in DMEM medium containing 10% fetal calf serum at a density of 4 x 10^6^ cells in a tissue culture-treated 10cm dish. The next day, 12μg of pMT123 DNA (pHA::ubiquitin) was transfected into cells using Lipofectamine 2000, following manufacturer’s procedure (Life Sciences). The mammalian HA::ubiquitin expression plasmid pMT123 was a kind gift from D. Bohmann and S. Lippard^60, 74^. The next day, transfection medium was removed and replaced with DMEM with 10μM MG132 (Calbiochem) at ∼30-60min prior to challenge with *L. pneumophila* at an MOI=10 for the indicated times ^75, 76^. Post infection, cells were lifted and washed in PBS, centrifuged at 4°C at 200 x g, and then cell pellets were immediately frozen on liquid nitrogen and stored at -80°C until used^22^.

For immunoprecipitations, anti-Rtn4 resin was generated by disuccinimidyl suberate crosslinking with anti-Rtn4 (Nogo N-18, Santa Cruz sc-11027) to Protein A/G Plus™ resin at 2μg/μl of packed resin as described in manufacturer protocol (Pierce crosslink IP kit). Samples were lysed in 1% Triton X-100 (20 mM Tris [pH 7], 150 mM NaCl, 5 mM MgCl_2_) containing protease inhibitor (Roche, Cat# 11697498001) with end-over-end rotation for 20 min at 4°C. Cleared cell lysates were obtained by centrifugation at ≥14,000 x g for 10 min at 4°C, diluted with an equal volume of detergent-free buffer, and mixed with washed resin (50% slurry). The mixture was incubated at 4°C via end-over-end rotation for ≥ 4 hr, then washed ≥ 5X in the buffer containing 0.1% Triton X-100 at 4°C.

For elution, Rtn4 resin in spin columns was incubated with 0.1M glycine (pH = 2.8) with 0.2% Triton X-100 for 5min, centrifuged, then eluate was reapplied to resin for additional 5 min incubation at room temperature, centrifuged and neutralized with 0.5M Tris (pH=10.55)^22^. Eluate bound to resin was resuspended with reducing SDS sample buffer, boiled, then fractionated by SDS-PAGE coupled with immunoblot analyses.

### Protein purification

Full-length *sdeC* genes were amplified by PCR and cloned into the bacterial expression vector pQE-80L (Qiagen) with *BamHI* and *SacI* restriction sites to generate a 6xHis-SdeC-StrepII fusion construct (Supplemental Table S1)^22^. Purified aliquots were stored at-80°C in 10% glycerol to minimize or eliminate freeze thaw cycle degradation.

### Deubiquitinase assay in presence of Sde modification

His-tagged recombinant SdeC and its derivatives were bound to Ni-NTA (Thermo Fisher Scientific, Cat# 88221) resin at 1 μg enzyme per 25μl of packed resin in 1X ART buffer (20mM Tris 10mM NaCl, pH7.4) for 1 hr at 4°C with end-over-end rotation. The beads were washed 3X with 1X ART buffer to remove non-adsorbed enzyme with microcentrifuge spin filters spun at 3000 x g at room temperature. After the final wash, each 25μl of pelleted resin was incubated with 2μg of recombinant human K63-linked (R&D systems, Cat# UC-320) or K48-linked (R&D Systems, Cat# UC-220) poly-ubiquitin and 500μM ε-NAD in 300 μl of 1X ART buffer for 2 hrs with end-over-end gentle rotation at 37°C. After incubation, the SdeC or its derivatives-bound resin were removed with microcentrifuge spin column filters, and the polyUb reaction mixture was collected. This polyUb fraction was then incubated with either 1) 100 nM human recombinant PolyHistidine-otubain 1( R&D systems, Cat# E-522B), 2) 50 nM recombinant USP2 catalytic domain (R&D systems, Cat# E-504), 3) 50 nM recombinant PolyHistidine-CYLD (R&D systems, Cat# E-556), or 50nM recombinant SdeC DUB (1-192) at 37°C in 1X ART buffer for 2 hr. A fraction of the reaction was removed, terminated by the addition of reducing sample buffer, and then heated to 50-55°C for 20-30min to avoid Ub chain aggregation. Relative DUB activity was determined by immunoblot probing of SDS-SDS-PAGE fractionated proteins using Image Studio software (LI-COR Biosciences) after 2 hr incubation by comparing the efficiency of cleavage of modified Ub relative to unmodified.

### Immunoblotting

Immunoblots were performed as described ^77^. Samples were fractionated by SDS-PAGE and transferred to nitrocellulose membranes. The membrane was blocked in 50 mM Tris-buffered saline with 0.1% Tween-20 (TBST, pH 8.0) containing 4% nonfat milk for 1 hr at room temperature and probed with primary antibodies: 1) anti-HA-probe (Santa Cruz Biotechnology, Cat# sc-7392, 1:500), 2) anti-Ub (FK1; Santa Cruz Biotechnology, Cat# sc-8017) or anti-actin (Sigma-Aldrich, Cat# A2066, 1:1,000) in TBST containing 2% BSA at 4°C overnight. After washing 3X with TBST, the membranes were incubated with secondary antibodies: donkey IRDye 680RD anti-mouse (LI-COR Biosciences, Cat# 926-68072, 1:20,000), Goat IRDye 680RD anti-rabbit (LI-COR Biosciences, Cat# 926-68071, 1:20,000), and/or donkey IRDye800CW anti-mouse (LI-COR Biosciences, Cat# 926-32212, 1:20,000), goat IRDye 800CW α-rabbit (LI-COR Biosciences, Cat# 926-32211, 1:20,000) for 45 min at room temperature. Capture and analysis were performed using Li-Cor Odyssey CLX scanner and Image Studio software (LI-COR Biosciences).

### Bead-binding assay for adaptors association with polyUb post Sde-mediated modification of Ub

His-tagged recombinant full-length SdeC derivatives were adsorbed to Ni-NTA (Thermo Fisher Scientific, Cat# 88221) resin at 1μg enzyme per 25 μl of packed resin in 1X ART buffer for 1 hr at 4°C with end-over-end rotation. The SdeC adsorbed resin was washed 3X with 1X ART buffer to remove nonadsorbed enzyme with microcentrifuge spin filters centrifuged at 3000 x g at room temperature. After the final SdeC resin wash, each 25 μl pelleted resin was resuspended with 4μg of recombinant human homotypic-linked polyubiquitin chains Ub_3-7_ (K63 or K48, 4μg/600μl reaction volume, Boston Biochem, R&D) and 500μM ε-NAD in 1X ART buffer for 2 hrs with gentle end-over-end rotation at 37°C. During incubation of SdeC and polyUb chains, 1μg recombinant PolyHis-p62 was adsorbed to 5 μl pre-equilibrated packed Ni-NTA resin (Thermo Fisher Scientific, Cat# 88221) via 1 hr, end-over-end rotation at 4°C in low adhesion 1.5 ml microcentrifuge tubes. Polyubiquitin chains were then separated from SdeC by collection of SdeC resin in a microcentrifuge column spin filter and then the polyUb fraction was incubated with recombinant PolyHis-p62 on Ni-NTA resin for 1 hr at room temperature with end-over-end rotation. The resin was washed 3-5X with 50mM Tris 0.1% BSA (vol/vol) pH7.5 at room temperature, then resin having adsorbed proteins and bound polyUb chains were eluted from the Ni-NTA beads with 250 mM imidazole diluted in 50 mM Tris 0.1% BSA (vol/vol) pH7.5. Samples were then analyzed by SDS-PAGE followed by immunoblot.

### Preparation of Sde-modified ubiquitin derivatives

For modification of Ub, 100 μg of K63-linked Ub tetramer (R&D Systems, Cat# UCB- 310 or South Bay Bio, SBB-UP0073) or monomeric Ub (R&D Systems, Cat# U-100H) were incubated together with 100 μM β-NAD and 20 nM SdeC variants (WT, C118S/DUB^-^, E859A/ART^-^ and H416A/PDE^-^) in 250 μl of 1X ART buffer at 37°C for 1 h. For DUB inhibitor pretreatment, 1 μM ubiquitin-propargylamide (R&D Systems, Cat# U-214) was incubated with 100 nM SdeC variants at room temperature for 30 min before adding to Ub_4_ (final concentration, 20 nM SdeC in 1X ART buffer). To remove the SdeC, 10 μl of a 50% slurry of Ni-NTA agarose resin (Thermo Fisher Scientific, Cat# 88221) was washed three times with 1X ART buffer, then added to the Ub-SdeC reaction for another 1hr with end-over-end rotation at 37°C. For removal of SdeC enzyme, the samples were centrifuged at 1000 x g for 2 min in a microcentrifuge spin filter column. The supernatant was then desalted using a NAP-5 column (Thermo Fisher Scientific, Cat# 45-000-151) and lyophilized with a Labconco Freezone12. Samples were resuspended in Ultrapure water to a final protein concentration of 1 mg/ml.

### LC-MS analysis of modified Ub_4_

Reversed-phase chromatography and mass spectrometry was performed on an Agilent 1260 Infinity LC system in line with an Agilent 6530 Q-TOF. Intact Ub_4_ samples were diluted in water and injected onto a Zorbax 300 SB-C8 Rapid Resolution HD 1.8 μm column (2.1 x 100 mm, Agilent) and were eluted with a water:acetonitrile gradient mobile phase with 0.1% formic acid (0.400 mL/min; 95% - 20% over 26 min). The mass spectrometer was utilized in positive mode with a dual electrospray ionization (ESI) source. MS spectra were acquired using the following settings: ESI capillary voltage, 4500 V; fragmentor, 250 V; gas temperature, 325°C; gas rate, 12.5 L/min; nebulizer, 50 psig. Data was acquired at rate of 5 spectra per second and scan range of 100 – 3000 m/z.

For trypsin digestion, Ub_4_ samples were incubated at 95 degrees in 1:1 water:t-butanol for 35 min, then allowed to cool to room temperature. DTT was added to a concentration of 3 mM, trypsin was added in a ratio of 25:1 Ub_4_:trypsin and incubated overnight. The reaction was then diluted in dH_2_O for mass spectrometry. Following tryptic digestion, peptide fragments were injected onto an AdvanceBio Peptide 2.7 μm column (2.1 x 150 mm, Agilent) and were eluted with a water:acetonitrile gradient mobile phase with 0.1% formic acid (0.400 ml/min; 95% - 5% over 19 min). MS spectra were acquired using the following settings: ESI capillary voltage, 4000 V; fragmenter, 150 V; gas temperature, 325°C; gas rate, 12 L/min; nebulizer, 40 psig. Data were acquired at a rate = 5 spectra per second and scan range of 300 – 3000 m/z. MS/MS spectra were acquired using the following settings: ESI capillary voltage, 4000 V; fragmentor, 150 V; gas temperature, 325°C; gas rate 12 l/min; nebulizer, 40 psig. MS/MS was acquired at 2 spectra per second at a mass range of 100 – 3000 m/z, with stringency set to a medium isolation width. After identification, precursor ions were subjected to iterative rounds of collision-induced dissociation in the collision chamber and subsequent mass identification. A ramped collision energy was used with a slope of 3.6 and offset of -4.8 as well as a slope of 3 and offset of 2.

Following acquisition, analysis was completed using multiple programs. Agilent MassHunter Qualitative Analysis (v. 7.00) was used to generate mass spectra, and extracted ion chromatograms (EIC). Agilent MassHunter Bioconfirm (v. B.010.00) was used to identify and match tryptic fragments of the Ub_4_ and pR modified Ub_4_ proteins, and identify b and y ion fragmentation from combined MS/MS spectra. For the control mono-Ub incubated in the presence or absence of SdeC, n=1 for each treatment. For Ub_4_ in absence of SdeC, n=2 biological replicates, from two different suppliers. For Ub_4_ incubated with SdeC, n=2 technical replicates for intact protein and n=7 technical replicates for trypsin digested protein. MS data were deposited in ProteomeXchange (http://www.proteomexchange.org/), accession number MSV000094791.

### SPR analysis of p62 binding to ubiquitin

The binding kinetics between p62 and modified Ub_4_ was determined by surface plasmon resonance (SPR) using a Biacore T200 system, with a runner buffer of HEPES Buffered Saline (HBS; 10 mM HEPES (pH =7.4), 150 mM NaCl) containing 30 µM of EDTA. 6xHis-tagged p62 was captured onto a Series S NTA sensor chip following manufacturer’s recommendations (Cytiva, Cat# 28994951). Briefly, 0.5 mM NiCl_2_ solution was injected at a flow rate of 10 µl/min for 60 seconds onto the experimental flow cell, followed by the injection of His-p62 (200 nM) at 5 ul/min for 420 seconds. Ub_4_ at various concentrations, that had been previously treated with SdeC derivatives, was injected into the flow cells at a rate of 30 µl/min for 180 seconds, followed by a 300 seconds dissociation phase. Regeneration of the surface was performed by two injections of 350 mM EDTA (30 ul/min, 60 seconds) followed by an injection of 6M guanidine hydrochloride with 50 mM NaCl (30 ul/min, 30 seconds). The binding kinetics were determined with Biacore Evaluation software (v3) using double referenced sensorgrams and 1:1 binding for the fitting model.

### Immunofluorescence of p62 at the LCV

BMDMs were plated on glass coverslips at 1-2 x10^5^/well in 24-well plates. Cells were challenged at a multiplicity of infection (MOI) = 2, centrifuged at 200 x g for 5 min, and incubated for the indicated time points. For 180 min infections, the cells were washed 3X in PBS at 60 min post-infection to remove non-internalized bacteria, then fresh culture media was added. At the infection conclusion, all cells were washed 3X in PBS, fixed in PBS containing 4% (wt/vol) paraformaldehyde, followed by permeabilization with ice-cold 100% methanol for 20 seconds. Coverslips were then washed 3X in PBS and blocked in PBS containing 2-4% BSA for at least 30 min. Cells were then stained in PBS with rat anti-*L. pneumophila* and rabbit anti- p62/SQSTM (Abcam, Cat# 109012, 1:400) for 1 hr and detected with donkey anti-rat Alexa Fluor 594 (Jackson ImmunoResearch, Cat# 712-585-153, 1:500) and goat anti-rabbit Alexa Fluor 488 (Jackson ImmunoResearch, Cat# 111-545-003, 1:500). Hoechst 33342 was used to label DNA.

### Image analysis

Representative images were taken using Zeiss Axiovert or Nikon inverted phase fluorescence microscopes, and image enhanced in Photoshop™ (Adobe), using linear corrections. For quantification of total polyUb intensity, Volocity software (PerkinElmer) was used. ROIs (regions of interest, 14 x 12 μm; oval) were set around LCVs, and pixel intensities were measured and subtracted by background intensity. For quantification of total p62 intensity, we set ROIs (0.3 x 0.42 inches; oval) around LCVs. Mean p62 intensity/pixel was measured in each ROI and subtracted by background intensity/pixel using Fiji image analysis software^78^.

## Data Availability Statement

Data and images that support the findings of this study are available in Source Data File. Mass Spectrometry data were deposited in ProteomeXchange (http://www.proteomexchange.org/), accession number MSV000094791. Source data are provided with this paper.

## Statement on Samples

All measurements were taken from distinct samples. The following panels analyzed data from the same set of samples: 5C and 5D; 5E and 5F; 5G and 5H; 5J and 5K; 6A and 6B. Panel 5L is a summary of panels in Fig. 5, as noted in Figure Legend.

## Supporting information

Complete Supplementary Tables

## Acknowledgements

KMK was supported by NIH training grant T32GM007310. This work was supported by NIAID grants R01AI113211 and R01AI146245 to RRI and NIGMS grant R01GM132422 to RAS. Panels 4A and 4B were used under terms of Creative Commons. We thank Drs Kevin Manera for review of the manuscript and Dr. Yuxin Mao for communication of results before publication.

## Author Contributions

KMK and MZ designed, performed and analyzed experiments as well as contributed to writing of the manuscript. SK performed and analyzed experiments. MSM designed, performed and analyzed experiments and contributed to writing of manuscript. ARC designed, performed and analyzed experiments as well as contributed to writing of manuscript. AT performed and analyzed experiments. RAS directed research, designed and analyzed experiments and contributed to writing of manuscript. RRI directed research, designed and analyzed experiments and contributed to writing of manuscript.

## Figure Legends

**Supplemental Table S1.** *Legionella pneumophila* strains used in this study.

**Supplemental Table S2.** *Escherichia coli* strains used in this study

**Supplemental Table S3.** Oligonucleotides used in this study.

**Supplemental Figures S1-S5**

