## Supplementary Fig. 1 for "Sde Proteins Coordinate Ubiquitin Utilization and Phosphoribosylation to Establish and Maintain the *Legionella* Replication Vacuole"

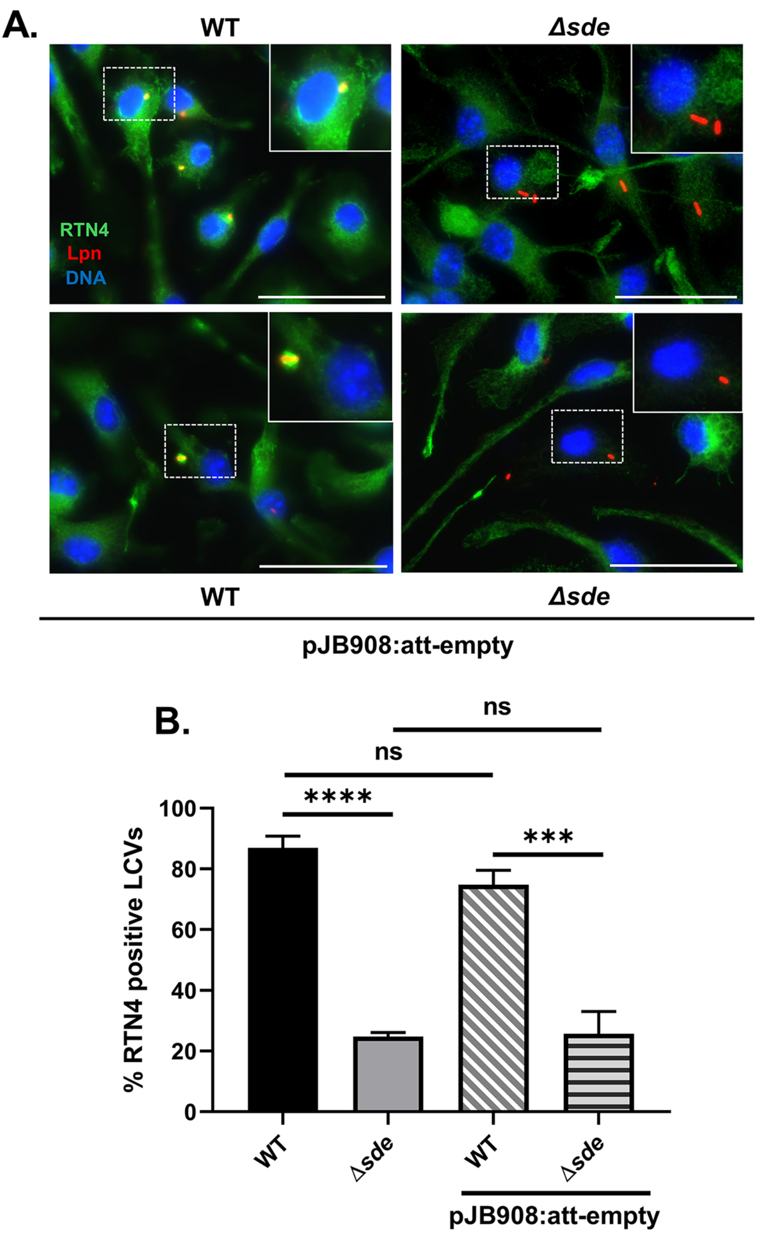


Fig. S1. The presence of empty vector has no effect on relative Rtn4 recruitment. (Linked to Fig. 1). Representative image of Rtn4 recruitment around the LCV by WT and Δsde mutant (scale bar= 5 μm). BMDMs were challenged at MOI =1 with WT, Δsde, WT/pJB908:att-empty, and Δsde/ pJB908:att-empty for 1 hr, fixed, permeabilized, and probed with anti-L. pneumophila (Alexa Fluor 594 secondary, red), anti-Rtn4 (Alexa Fluor 488 secondary, green), and Hoechst (nucleus, blue). (B) Quantification of Rtn4-positive LCVs. Images were captured and over 250 LCVs were analyzed per coverslip, and data were pooled from three independently performed experiments. Data are represented as mean ± SEM. Statistics performed by one-way ANOVA with Dunnett's multiple comparisons; ns= non-significant, *p<0.05, **p<0.01, ***p<0.001, ****p < 0.0001)
