## Supplementary FIg. 2 for "Sde Proteins Coordinate Ubiquitin Utilization and Phosphoribosylation to Establish and Maintain the *Legionella* Replication Vacuole"

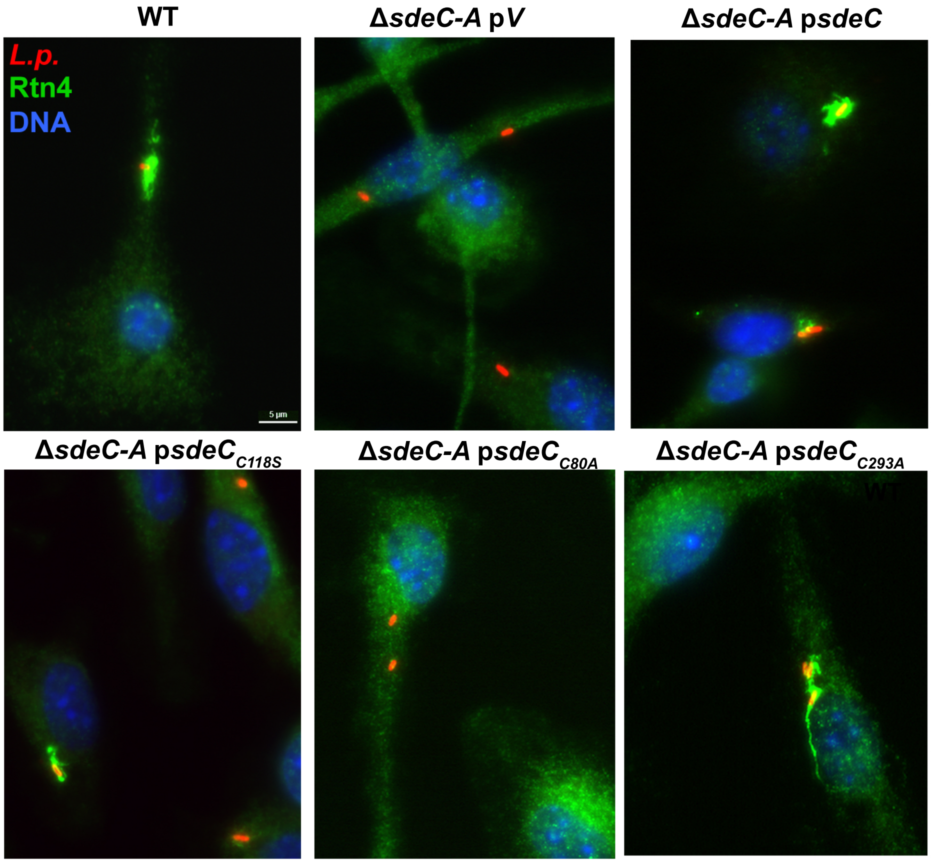


**Fig. S2. Sde DUB domain is required for efficient phosphoribose-ubiquitination of Rtn4.** Representative images of Rtn4 recruitment, scale bar 5 µm. BMDMs were challenged with *L. pneumophila* strains for 1 hr, fixed, permeabilized and probed as described in Fig. 1. Strains used are WT, Lp02; Δ*sdeC-A*, Δ*lpg2153-2157* (KK034); Δ*sdeC-A* psdeC, Δ*sdeC-A* expressing SdeC; Δ*sdeC-A* p*sdeC*$_{C118S}$, Δ*sdeC-A* expressing SdeC DUB mutant (C118S); Δ*sdeC-A* p*sdeC*$_{D80A}$, Δ*sdeC-A* expressing SdeC DUB mutant (D80A); Δ*sdeC-A* p*sdeC*$_{C293A}$, Δ*sdeC-A* expressing SdeC DUB mutant (C293A).
