## Supplementary Fig. 3 for "Sde Proteins Coordinate Ubiquitin Utilization and Phosphoribosylation to Establish and Maintain the *Legionella* Replication Vacuole"

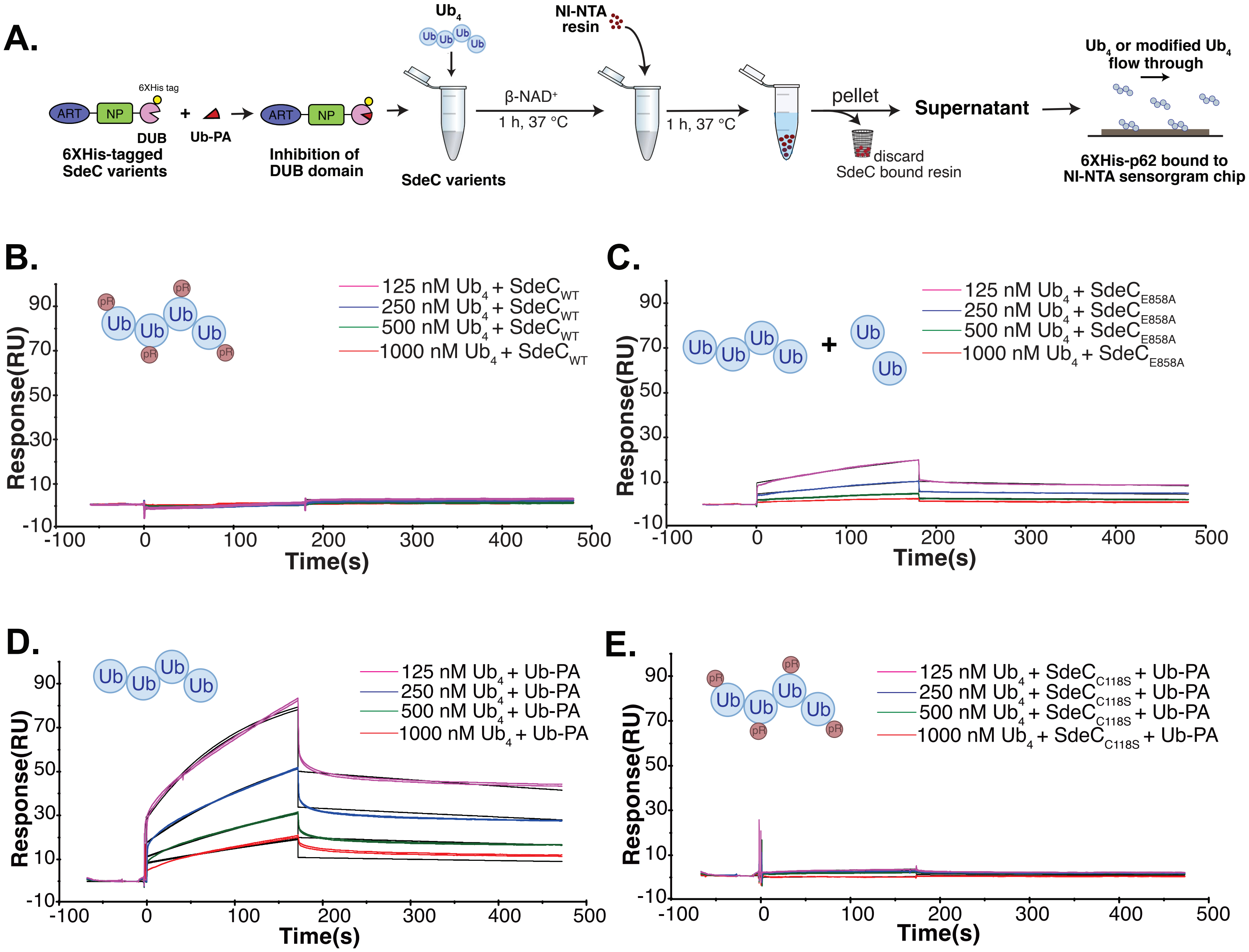


**Figure S3. The presence of a DUB inhibitor allows analyss of binding to p62.** A. SdeC variants were incubated with tetra-Ub in the presence or absence of DUB inhibitor Ub-PA and analyzed for binding to p62 using SPR analysis. B-E analysis of p62 binding after SdeC treatment of $Ub_4$ in presence or absence of Ub-PA.
