## Supplementary Fig. 4 for "Sde Proteins Coordinate Ubiquitin Utilization and Phosphoribosylation to Establish and Maintain the *Legionella* Replication Vacuole"

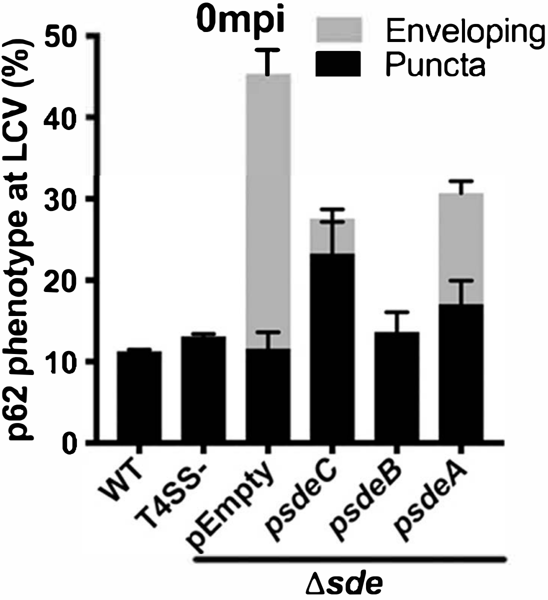


**Fig. S4.** Multiple morphological forms of p62 are associated with the LCV (Linked to Fig. 4). Data shown are from the identical coverslips as displayed in Fig. 4C, with quantification of enveloped LCVs shown in Fig. 4C overlayed with quantification of puncta. Puncta are determined based on morphology displayed in Fig. 4A and acquired at same time as acquired for Fig. 4C. Puncta association was indistinguishable for WT, T4SS- and Δsde strains.
