## Supplementary Fig. 5 for "Sde Proteins Coordinate Ubiquitin Utilization and Phosphoribosylation to Establish and Maintain the *Legionella* Replication Vacuole"

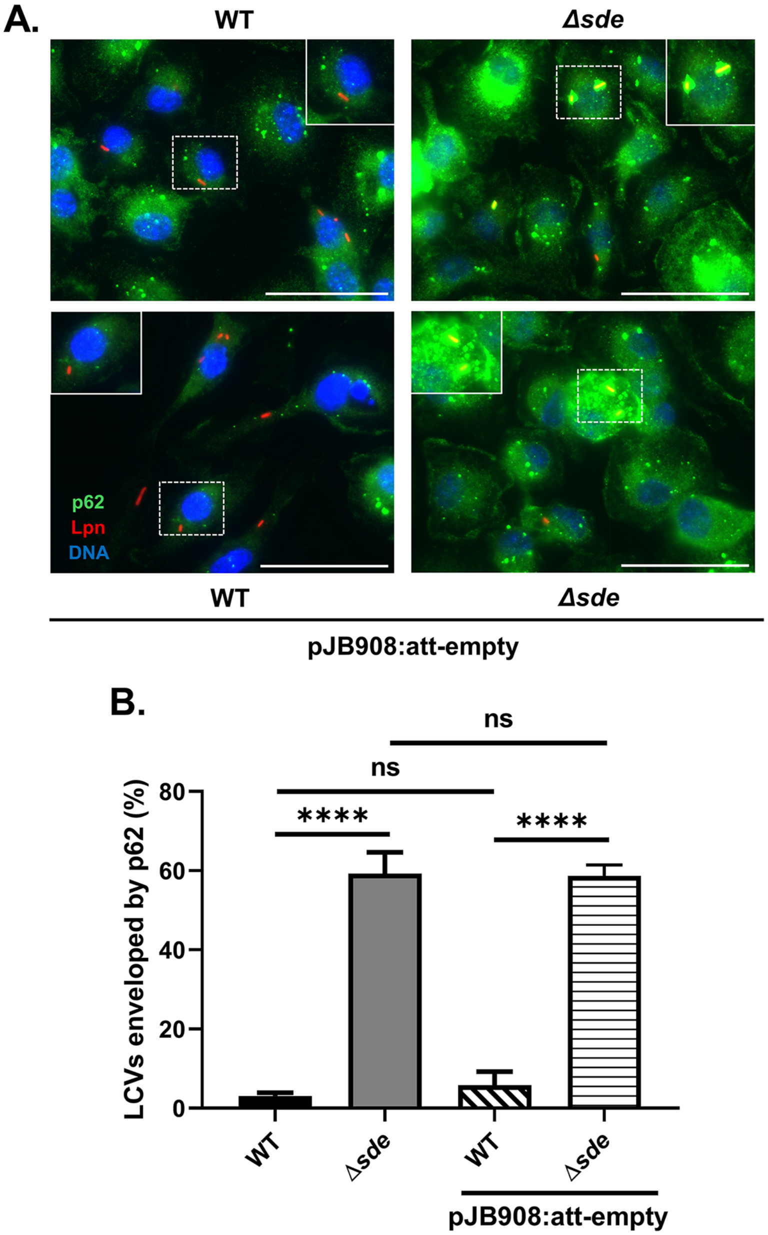


**Fig. S5.** The presence of empty vector has no effect on relative p62 recruitment. (Linked to Figs. 5,6). A representative image of p62/SQSTM1 recruitment around the LCV (scale bar= 5 μm). BMDMs were challenged at MOI =2 for 20 minutes, fixed, permeabilized, and probed with anti-L. pneumophila (Alexa Fluor 594 secondary, red), anti-p62 (Alexa Fluor 488 secondary, green), and Hoechst (nucleus, blue). (B) Quantification of LCVs enveloped with p62/SQSTM1. Images were captured and over 150 LCVs were analyzed per coverslip, and data were pooled from three independently performed experiments. Data are represented as mean ± SEM. Statistic performed by one-way ANOVA with Dunnett's multiple comparisons; ns= non-significant, *$p<0.05$, **$p<0.01$, ***$p<0.001$, ****$p < 0.0001$).
