## Supplementary material for "Sde Proteins Coordinate Ubiquitin Utilization and Phosphoribosylation to Establish and Maintain the *Legionella* Replication Vacuole": Complete Supplementary Tables

**Supplemental Table S1. *Legionella pneumophila* strains used in this study.**

| **Strain** | **Genotype** | **Description** | **Reference** |
| --- | --- | --- | --- |
| Lp02 | (WT) Philadelphia 1, *thyA rpsL hsdR* | Wild type | ^1^ |
| Lp03 | Lp02 *dotA03* | Translocation deficient | ^1^ |
| Lp02 pvector | Lp02 (pJB908att-empty) | pJB908 Gateway™ vector *ΔccdB* (PolyHis/c-myc epitope tag vector) | ^2^ |
| Lp03 pvector | Lp03 (pJB908att-empty) | pJB908 Gateway™ vector *ΔccdB* (PolyHis/c-myc epitope tag vector) | ^2^ |
| KK034 | Lp02*ΔsdeC-sdeA* *(Δlpg2153-2157)* *kan^R^ P_ahpC_*::*lux* | *sdeC-sdeA* deletion, Lux^+^ | ^2^ |
| KK034 pSdeC_WT_ | KK034 (pJB908att-*sdeC_WT_*) | PolyHis/c-myc::SdeC WT | ^2^ |
| KK034 pSdeC_C118S_ | KK034 (pJB908att-*sdeC_C118S_*) | PolyHis/c-myc::SdeC DUB- | ^2^ |
| KK034 pSdeB_WT_ | KK034 (pJB908att-*sdeB_WT_*) | PolyHis/c-myc::SdeB WT | ^2^ |
| KK034 pSdeA_WT_ | KK034 (pJB908att-*sdeA_WT_*) | PolyHis/c-myc::SdeA WT | ^2^ |
| JV6113 | Lp02 *ΔsidE ΔsdeC ΔsdeBA* (*Δlpg0234, Δlpg2153 Δlpg2156-2157*) | *sidE* family deletion | ^3^ |
| KK099 | JV6113 *kan^R^ P_ahpC_*::*lux* | *sidE* family deletion, Lux+ | ^2^ |
| KK099 pvector | *ΔsidE ΔsdeC ΔsdeBA* (pJB908att-empty) | pJB908 Gateway™ vector *ΔccdB* (PolyHis/c-myc epitope tag vector) | ^2^ |
| KK099 pSdeC_WT_ | KK099 (pJB908att-*sdeC_WT_*) | PolyHis/c-myc::SdeC WT | ^2^ |
| KK099 pSdeC_C118S_ | KK099 (pJB908att-*sdeC_C118S_*) | PolyHis/c-myc::SdeC DUB- | ^2^ |
| KK099 pSdeC_H416A_ | KK099 (pJB908att-*sdeC_H416A_*) | PolyHis/c-myc::SdeC NP- | ^2^ |
| KK099 pSdeC_E859A_ | KK099 (pJB908att-*sdeC_E858A_*) | PolyHis/c-myc::SdeC ART- | ^2^ |
| KK099 pSdeC_C118S/E859A_ | KK099 (pJB908att-*sdeC_E858A_*) | PolyHis/c-myc::SdeC  DUB-/ART- | ^2^ |
| KK099 pSdeC_C118S/H416A_ | KK099 (pJB908att-*sdeC_E858A_*) | PolyHis/c-myc::SdeC  DUB-/NP- | ^2^ |
| KK099 pSdeB_WT_ | KK099 (pJB908att-*sdeB_WT_*) | PolyHis/c-myc::SdeB WT | ^2^ |
| KK099 pSdeB_C118S_ | KK099 (pJB908att-*sdeC_R763A_*) | PolyHis/c-myc::SdeB DUB- | ^2^ |
| KK099 pSdeB_E859A_ | KK099 (pJB908att-*sdeB_E859A_*) | PolyHis/c-myc::SdeB ART- | ^2^ |
| KK099 pSdeB_H416A_ | KK099 (pJB908att-*sdeB_E859A_*) | PolyHis/c-myc::SdeB NP- | ^2^ |
| KK099 pSdeB_C118S/E859A_ | KK099 (pJB908att-*sdeC_E858A_*) | PolyHis/c-myc::SdeB  DUB-/ART- | ^2^ |
| KK099 pSdeB_C118S/H416A_ | KK099 (pJB908att-*sdeC_E858A_*) | PolyHis/c-myc::SdeB  DUB-/NP- | ^2^ |
| KK099 pSdeA_WT_ | KK099 (pJB908att-*sdeA_WT_*) | PolyHis/c-myc::SdeA WT | ^2^ |
| KK099 pSdeA_C118S_ | KK099 (pJB908att-*sdeA_R766A_*) | PolyHis/c-myc::SdeA DUB- | ^2^ |

**Supplemental Table S2. *Escherichia coli* strains used in this study.**

| ***E. coli*** | | |
| --- | --- | --- |
| BL21 DE3 | *F^-^ ompT hsdSB gal dcm (DE3)* |  |
| DH5α | *supE44 ΔlacU169(Φ80lacZDM15) hsdR17 recA1 endA1 gyrA96 thi-1 relA1* |  |
| MT607 pRK600 | *recA56 pro-82 thi-1 hsdR17 supE44* (RK600 XL-1) | ^4^ |

**Supplemental Table S3. Oligonucleotides used in this study.**

| **Primers** | |
| --- | --- |
| **Description** | **Sequence (5’ 🡪 3’)** |
| SdeC flanking primer with *attB1* | GGGGACAAGTTTGTACAAAAAAGCAGGCTTCATGCCTAAATACGTAGAAGGGGTAG |
| SdeC flanking primer with *attB2* | GGGGACCACTTTGTACAAGAAAGCTGGGTCCTATTAGAAACCATACCTTATGTCATCG |
| SdeB flanking primer with *attB1* | GGGGACAAGTTTGTACAAAAAAGCAGGCTTAATGCCTAAATATGTAGAAGGGGTAG |
| SdeB flanking primer with *attB2* | GGGGACCACTTTGTACAAGAAAGCTGGGTCCTATTAAAAGTAACCACTTTCTCGGTG |
| SdeA flanking primer with *attB1* | GGGGACAAGTTTGTACAAAAAAGCAGGCTTAATGCCTAAGTATGTCGAAGGGGTAG |
| SdeA flanking primer with *attB2* | GGGGACCACTTTGTACAAGAAAGCTGGGTCCTATTAAAATCCTATAGTTTTTTTATTGG |
| SdeC – pQE-80L | GGGGGATCCATGCCTAAATACGTAGAAGGG |
| SdeC – pQE-80L (C-terminus StrepII) | GGGGAGCTCTTATTATTTTTCGAACTGCGGGTGGCTCCAAGCGCTGAAACCATACC |
| Quickchange: SdeC C118S | GAGCCCGGCCTAAACAAAGGCCTATCTGGCTATTGGGTAGCCTC |
| Quickchange: SdeC C118S | GAGGCTACCCAATAGCCAGATAGGCCTTTGTTTAGGCCGGGCTC |
| Quickchange: SdeB C118S | GAAGCAACCCAATAGCCAGATAGCCCCTTGTTTAGGC |
| Quickchange: SdeB C118S | GGCCTAAACAAGGGGCTATCTGGCTATTGGGTTGCTTC |
| Quickchange: SdeA C118S | AGGCGACCCAGTAGCCGGATAATCCCATATTCAAACC |
| Quickchange: SdeA C118S | GGTTTGAATATGGGATTATCCGGCTACTGGGTCGCCTC |
| Quickchange: SdeC H416A | CAT ATC CTG ATA AAT CAA GGC GCT ATG GTC GAT TTA ATG AGG AC |
| Quickchange: SdeC H416A | GT CCT CAT TAA ATC GAC CAT AGC GCC TTG ATT TAT CAG GAT ATG |
| Quickchange: SdeC/SdeB R763A | CTCCGAAAAAACTGTATGCTGGATTAAATCTGCCACAAG |
| Quickchange: SdeC/SdeB R763A | CTTGTGGCAGATTTAATCCAGCATACAGTTTTTTCGGAGC |
| Quickchange: SdeC E859A | CATATGGCTGGTTCCGAAGATGCATTTTCCGTTTATTTGC |
| Quickchange: SdeC E859A | GCAAATAAACGGAAAATGCATCTTCGGAACCAGCCATATG |
| Quickchange: SdeB E859A | CATATGACTGGATCAGAAGATGCATTTTCCGTTTATTTGCC |
| Quickchange: SdeB E859A | CAAATAAACGGAAAATGCATCTTCTGATCCAGTCATATGG |

**Table 3. Plasmids used in this study.**

| **Plasmids** |  | **Ref** |
| --- | --- | --- |
| Transfection screen |  | Table 2.5. |
| pQE-80L | Expression vector for N-terminal PolyHis tagged proteins | Qiagen |
| pQE-SdeC^WT^ | pQE-80L *sdeC*, C-terminus StrepII tag | ^2^ |
| pQE-SdeA^WT^ | pQE-80L *sdeA*, C-terminus StrepII tag | ^2^ |
| pQE-SdeC^C118S^ | pQE-80L *sdeC* C118S, C-terminus StrepII tag | ^2^ |
| pQE-SdeC^H286A^ | pQE-80L *sdeC* H286A, C-terminus StrepII tag | ^2^ |
| pQE-SdeC^R342A^ | pQE-80L *sdeC* R342A, C-terminus StrepII tag | ^2^ |
| pQE-SdeC^H416A^ | pQE-80L *sdeC* H416A, C-terminus StrepII tag | ^2^ |
| pQE-SdeC^R422A^ | pQE-80L *sdeC* R422A, C-terminus StrepII tag | ^2^ |
| pQE-SdeC^R763A^ | pQE-80L *sdeC* R763A, C-terminus StrepII tag | ^2^ |
| pQE-SdeC^E859A^ | pQE-80L *sdeC* E859A, C-terminus StrepII tag | ^2^ |
| pSR47-*P_ahpC_::lux* | pSR47 containing *P. luminescens* *lux* operon | ^5^ |
| pSR47s | *oriT*RP4 *ori*R6K *kan sacB* | ^4^ |
| pJB2199 | SR47S Δ*sdeC-sdeA* (contains region flanking *sdeA* and *sdeC*) | ^3^ |
| pJB908 | RSF1010 *thyA*^+^ *bla^+^ Δmob* | ^6^ |
| pJB908attR | pJB908 PolyHis/c-myc-*attR1*-[Cm^R^ *ccdB*]*-attR2* | ^2^ |
| pJB908att-empty | pJB908 PolyHis/c-myc-*attB1-attB2* | ^2^ |
| pSdeC_WT_ | pJB908 PolyHis/c-myc-*attB1-sdeC_WT_-attB2* | ^2^ |
| pSdeC_C118S_ | pJB908 PolyHis/c-myc-*attB1-sdeC_C118S_-attB2* | ^2^ |
| pSdeC_C293A_ | pJB908 PolyHis/c-myc-*attB1-sdeC_C293A_-attB2* | ^2^ |
| pSdeC_H416A_ | pJB908 PolyHis/c-myc-*attB1-sdeC_H416A_-attB2* | ^2^ |
| pSdeC_E859A_ | pJB908 PolyHis/c-myc-*attB1-sdeC_E859A_-attB2* | ^2^ |
| pSdeC_C118S/E859A_ | pJB908 PolyHis/c-myc-*attB1*-*sdeC_C118S/E859A_-attB2* | ^2^ |
| pSdeB_C118S/H416A_ | pJB908 PolyHis/c-myc-*attB1*-*sdeB_C118S/H416A_-attB2* | ^2^ |
| pSdeB_WT_ | pJB908 PolyHis/c-myc-*attB1-sdeB_WT_-attB2* | ^2^ |
| pSdeB_C118S_ | pJB908 PolyHis/c-myc-*attB1-sdeB_C118S_-attB2* | This study |
| pSdeB_H416A_ | pJB908 PolyHis/c-myc-*attB1*-*sdeB_E859A_-attB2* | ^2^ |
| pSdeB_E859A_ | pJB908 PolyHis/c-myc-*attB1*-*sdeB_E859A_-attB2* | ^2^ |
| pSdeB_C118S/E859A_ | pJB908 PolyHis/c-myc-*attB1*-*sdeB_C118S/E859A_-attB2* | ^2^ |
| pSdeB_C118S/H416A_ | pJB908 PolyHis/c-myc-*attB1*-*sdeB_C118S/H416A_-attB2* | ^2^ |
| pSdeA_WT_ | pJB908 PolyHis/c-myc-*attB1*-*sdeA_WT_-attB2* | ^2^ |
| pSdeA_C118S_ | pJB908 PolyHis/c-myc-*attB1-sdeA_C118S_-attB2* | This study |

1	Berger, K. H. & Isberg, R. R. Two distinct defects in intracellular growth complemented by a single genetic locus in Legionella pneumophila. *Molecular microbiology* **7**, 7-19 (1993). <https://doi.org:10.1111/j.1365-2958.1993.tb01092.x>

2	Kotewicz, K. M. *et al.* A Single Legionella Effector Catalyzes a Multistep Ubiquitination Pathway to Rearrange Tubular Endoplasmic Reticulum for Replication. *Cell Host Microbe* **21**, 169-181 (2017). <https://doi.org:10.1016/j.chom.2016.12.007>

3	Jeong, K. C., Sexton, J. A. & Vogel, J. P. Spatiotemporal regulation of a Legionella pneumophila T4SS substrate by the metaeffector SidJ. *PLoS Pathog* **11**, e1004695 (2015). <https://doi.org:10.1371/journal.ppat.1004695>

4	Berger, K. H., Merriam, J. J. & Isberg, R. R. Altered intracellular targeting properties associated with mutations in the Legionella pneumophila dotA gene. *Molecular microbiology* **14**, 809-822 (1994). <https://doi.org:10.1111/j.1365-2958.1994.tb01317.x>

5	Coers, J., Vance, R. E., Fontana, M. F. & Dietrich, W. F. Restriction of Legionella pneumophila growth in macrophages requires the concerted action of cytokine and Naip5/Ipaf signalling pathways. *Cellular microbiology* **9**, 2344-2357 (2007). <https://doi.org:10.1111/j.1462-5822.2007.00963.x>

6	Bardill, J. P., Miller, J. L. & Vogel, J. P. IcmS-dependent translocation of SdeA into macrophages by the Legionella pneumophila type IV secretion system. *Molecular microbiology* **56**, 90-103 (2005). <https://doi.org:10.1111/j.1365-2958.2005.04539.x>
